# Evaluating chromatin accessibility differences across multiple primate species using a joint modelling approach

**DOI:** 10.1101/617951

**Authors:** Lee E. Edsall, Alejandro Berrio, William H. Majoros, Devjanee Swain-Lenz, Shauna Morrow, Yoichiro Shibata, Alexias Safi, Gregory A. Wray, Gregory E. Crawford, Andrew S. Allen

## Abstract

Changes in transcriptional regulation are thought to be a major contributor to the evolution of phenotypic traits, but the contribution of changes in chromatin accessibility to the evolution of gene expression remains almost entirely unknown. To address this important gap in knowledge, we developed a new method to identify DNase I Hypersensitive (DHS) sites with differential chromatin accessibility between species using a joint modeling approach. Our method overcomes several limitations inherent to conventional threshold-based pairwise comparisons that become increasingly apparent as the number of species analyzed rises. Our approach employs a single quantitative test which is more sensitive than existing pairwise methods. To illustrate, we applied our joint approach to DHS sites in fibroblast cells from five primates (human, chimpanzee, gorilla, orangutan, and rhesus macaque). We identified 89,744 DHS sites, of which 41% are identified as differential between species using the joint model compared with 33% using the conventional pairwise approach. The joint model provides a principled approach to distinguishing single from multiple chromatin accessibility changes among species. We found that non differential DHS sites are enriched for nucleotide conservation. Differential DHS sites with decreased chromatin accessibility relative to rhesus macaque occur more commonly near transcription start sites (TSS), while those with increased chromatin accessibility occur more commonly distal to TSS. Further, differential DHS sites near TSS are less cell type-specific than more distal regulatory elements. Taken together, these results point to distinct classes of DHS sites, each with distinct characteristics of selection, genomic location, and cell type specificity.

## Introduction

It has long been hypothesized that phenotypic differences between species are more often due to genetic variation in non-coding regulatory regions than in protein-coding regions (King and Wilson 1975; Wray 2007; Wittkopp and Kalay 2011). The development of diverse genome-wide assays, combined with the publication of primate reference genomes, has allowed identification of inter-species differences in gene expression (Cáceres *et al*. 2003; Gilad *et al*. 2006; Blekhman *et al*. 2008; Brawand *et al*. 2011), DNA methylation (Pai *et al*. 2011; Zeng *et al*. 2012; Hernando-Herraez *et al*. 2013), histone modifications (Zhou *et al*. 2014; Villar *et al*. 2015), transcription factor binding motifs (Dermitzakis and Clark 2002; Odom *et al*. 2007; Schmidt *et al*. 2010), chromatin accessibility (Shibata *et al*. 2012; Gallego Romero *et al*. 2018), and alternative splicing (Blekhman *et al*. 2010; Barbosa-Morais *et al*. 2012). These differences in molecular function among primate species can provide valuable insights into species-specific trait differences, including disease risk (Prabhakar *et al*. 2008; Boyd *et al*. 2015; Prescott *et al*. 2015).

Conventional approaches to analyzing comparative functional genomic data employ multiple pairwise comparisons to detect differences between species (Robinson *et al*. 2010; Love *et al*. 2014). While these approaches work well with a few species, certain limitations with pairwise comparisons become apparent as the number of analyzed species increases. First, the multiple comparisons burden imposed by species number scales exponentially, reducing sensitivity. This is not an issue for the majority of published studies, which consider two or three species, but it quickly becomes constraining with additional species. Second, pairwise comparisons only consider part of the overall data when assessing whether a significant difference exists between any two species. Joint consideration of all the available data can provide a more informed inference of true differences. Third, when analyzing data from more than two ingroups species, the possibility of multiple state changes arises. In such cases, pairwise comparisons rely on a somewhat *post hoc* approach to resolve the phylogenetic history.

To address these concerns, we introduce a negative binomial generalized linear model that jointly models chromatin accessibility data from all available species and replicates. Regardless of the number of ingroup species, this method requires only one test to determine whether a given open chromatin site is differential among species. In contrast, the conventional pairwise approach uses *n-1* tests for *n* species, requiring a Benjamini-Hochberg correction that is *n-1* times larger than our approach. We applied our joint model to chromatin accessibility DNase-seq data from cultured skin fibroblasts obtained from four great apes (human, chimpanzee, gorilla, and orangutan) and an outgroup (rhesus macaque). We demonstrate that the joint modeling approach mitigates some of the challenges that arise when applying a pairwise approach to multiple ingroup species.

To facilitate application of our joint modeling approach with other data sets, we created a GitHub repository (http://github.com/ledsall/2019primate) with the script used to identify and classify differential sites, along with instructions for necessary modifications. Although we use a single script to both identify and classify differential sites, the steps can be separated and combined with other methods for classification (e.g. phylogenetically based methods).

## Materials and Methods

### DNase-seq Experiments and Sequencing

Fibroblast cell lines from 15 individuals comprising 3 biological replicates from each of five primate species (human, chimpanzee, gorilla, orangutan, and rhesus macaque) were obtained from Coriell (**Supplementary Table 1**). It is estimated that human and chimpanzee diverged 7 million years ago; gorilla diverged from the human-chimpanzee ancestor 10 million years ago; orangutan diverged from the human-chimpanzee-gorilla ancestor 18 million years ago; and rhesus macaque diverged from the human-chimpanzee-gorilla-orangutan ancestor 30 million years ago (Schrago and Voloch 2013). DNase-seq experiments were performed as previously described (Shibata *et al*. 2012). DNase-seq libraries were generated from 50 million cells and sequenced on Illumina instruments (**Supplementary Table 1)**.

### DNase-seq Read Mapping and Conversion to Human Genome

Due to the use of MmeI to generate DNase-seq libraries (Boyle *et al*. 2008), genomic DNA fragments are only 20 bases long. Therefore, sequencing reads were trimmed to 20 bases using a custom perl script. Reads were mapped to the species’ native genome: hg19 for human, panTro4 for chimpanzee, gorGor3 for gorilla, ponAbe2 for orangutan, and rheMac3 for rhesus macaque (Lander *et al*. 2001; Chimpanzee Sequencing and Analysis Consortium 2005; Locke *et al*. 2011; Yan *et al*. 2011; Scally *et al*. 2012). Reads were mapped using Bowtie version 0.12.9 (Langmead *et al*. 2009) (parameters: -trim5 0 --trim3 0 -m 1 -l 20) as part of a custom two-step pipeline. In the first step (“tier 1”), reads were required to match to a unique location with no mismatches (parameter: -n 0). In the second step (“tier 2”), unmapped reads from step one were re-mapped with a relaxed mismatch parameter of one mismatch (parameter: -n 1). Reads that mapped to multiple locations or had more than one mismatch were discarded. Samtools version 0.1.19-44428cd (Li *et al*. 2009) was used to convert the sam files from each step to bam files, merge them into one file, and remove duplicate reads (defined as having the same chromosomal coordinates). Bedtools v2.17.0 (Quinlan and Hall 2010) was used to convert the bam files to bed files. Details on the number of reads in the input files and at each step are included in **Supplementary Table 2**.

Reads from the non-human samples were converted from their native genomic coordinates to hg19 coordinates using a three-step process that removed reads lacking a one-to-one relationship between the genomes. In each step, read coordinates were converted from one genome to the other using the UCSC liftOver software (Hinrichs *et al*. 2006) with a minMatch parameter of 0.8, which requires that 80% of the read maps to the new genome. Note that this parameter filters only on the presence or absence of DNA, not nucleotide identity. In the first step, read coordinates were converted from their native genome to hg19. Read coordinates that successfully lifted to hg19 were then lifted back to the native genome. Read coordinates that did not lift back to the same coordinates on the native genome were removed. Reads that did lift back to the same coordinates were lifted back to hg19 for further processing. An additional filtering step was added to ensure the reads were not part of a duplicated region. In that step, overlapping reads on the native genome were merged into a region, which was then lifted to hg19. Regions that failed to lift uniquely to hg19 were flagged and reads that overlapped them were removed. Because some of the samples were from males and some were from females (**Supplementary Table 1**), we removed reads that mapped or lifted over to the human X or Y chromosomes to eliminate any sex-specific bias. Details on the number of reads lifted over and remaining after removal of sex chromosomes are included in **Supplementary Table 2**.

Phylogenetic trees were drawn with ggtree (Yu *et al*., 2016).

### DHS Site Identification and Filters

To avoid bias due to large differences in depth of library sequencing, 20 million reads were randomly selected from samples with library sizes greater than 20 million reads to keep all libraries approximately the same size. The random sampling was performed after the conversion to the human genome and filtering steps.

First, we identified DHS sites in each sample by performing peak calling using the MACS2 callpeak command with an FDR cutoff of 5% (Zhang *et al*. 2008) (version 2.1.0.20150420; parameters: --nomodel --extsize 20 --qvalue .05). Next, we identified per-species DHS sites. For each species, we used bedtools v2.17.0 (Quinlan and Hall 2010) to identify DHS sites by taking the union set of peaks that were found in at least 2 out of 3 biological replicates. We used bedtools to remove DHS sites that overlapped the ENCODE blacklist (http://hgdownload.cse.ucsc.edu/goldenPath/hg19/encodeDCC/wgEncodeMapability/wgEncodeDacMapabilityConsensusExcludable.bed.gz) (Rosenbloom *et al*. 2013). Lastly, we generated the master set of DHS sites to use for cross-species comparisons by using bedtools to create the union set of DHS sites identified in each species then applying two filters. In the first filter, we removed DHS sites without at least 95% genomic coverage between the start and stop coordinates for each of the species. Genomic coverage was determined using the Multiz Alignment MAF file (http://hgdownload.cse.ucsc.edu/goldenPath/hg19/multiz100way/maf/) from the UCSC Genome Browser Multiz Alignment of 100 Vertebrates (Blanchette *et al*. 2004), the Galaxy MAF Coverage Stats tool at usegalaxy.org (Afgan *et al*. 2018), and a custom perl script (filter_regions_based_on_conservation_coverage.pl; available in the GitHub repository). Next, we assigned read counts to each DHS site using bedtools. In the second filter, we used a custom perl script (zero_count_filter_HCGOM_min_2_replicates.pl; available in the GitHub repository) to remove DHS sites without DNase-seq sequence reads in at least two biological replicates from each species because they may be indicative of regions that were not sequenced or cannot be uniquely aligned. In other words, we expect at least some level of background DNase cutting across the genome. See **Supplementary Table 3** for DHS site counts before and after each filtering step.

### Principal Components Analysis

We performed a principal components analysis on the read counts from the 15 samples **(Supplementary Figure 1)**. We first normalized the counts by library size then ran the R prcomp function with the center and scale parameters set to true. We also performed a principal components analysis on the read counts from the 15 samples plus 3 additional chimpanzee samples from Pizzollo *et al*. 2018 **(Supplementary Figure 1)**. See **Supplementary Table 1** for details on the samples.

### Differential Site Identification and Classification

The read counts for each DHS site were used as input to a custom R script (GO.step10.run_glm.R; available in the GitHub repository) that identified and classified differential DHS sites. To address the over-dispersion problem inherent in count-based sequencing data, we used the R package DSS (Wu *et al*. 2013) to calculate a dispersion parameter for each DHS site, as well as a normalization offset (based on total library size) for each sample. For each DHS site, the read counts, dispersion parameter, and normalization offset were fit using a negative binomial generalized linear model with a log link function. Specifically, we fit two models: a species informed model and a null model in which species was not predictive of normalized counts. The species informed model models the expected counts by *log* (*λ_j_*) = *α*_*j*_ + *β*_*m*_ + *x*_*j*_^*T*^*β*, where λ represents expected counts, j indexes the sample, *α_j_* is a normalization offset for sample j, and *β*_*m*_ represents the expected counts for rhesus macaque (and is analogous to an intercept parameter). The design vector ***x_j_*** indicates to which species the j^th^ sample belongs and *x*_*j*_^*T*^ denotes its transpose. This design vector has 4 elements comprised of indicator functions of whether the sample is human, chimpanzee, gorilla, or orangutan. Specifically, *x*_*j*_^*T*^ = (1 0 0 0) if the j^th^ sample is human; *x*_*j*_^*T*^ = (0 1 0 0) if the j^th^ sample is chimpanzee; *x*_*j*_^*T*^ = (0 0 1 0) if the j^th^ sample is gorilla; and *x*_*j*_^*T*^ = (0 0 0 1) if the j^th^ sample is orangutan. Because accessibility changes are relative to rhesus macaque, it is used as the intercept in the models and does not have an indicator function. The vector *β* = (*β_h_*, *β*_*c*_, *β_g_*, *β_o_*)^*T*^ parameterizes the change in expected counts between each species and rhesus macaque. The null model assumes the vector *β* is zero and models the expected counts by *log* (λ_*j*_) = *α*_*j*_ + *β*_*m*_. These models were fit using the R package glm using negative.binomial (from package MASS) as the family. The DSS normalization offset value was used for the offset parameter. The inverse of the DSS dispersion parameter value was used for the theta parameter in the negative.binomial family function. The difference in deviances between these two models was used to form a likelihood ratio test of whether the site was differential. A Benjamini-Hochberg correction was performed using the R function p.adjust. DHS sites with a corrected p-value of less than 0.01 were classified as differential.

To determine which species (or combination of species) were different from rhesus macaque, 15 contrasts were constructed using the *β* values estimated in the regression model detailed above. A *β* value represents the change in accessibility for that species compared to rhesus macaque and each contrast tests the *β* values for a different combination of species. Five contrasts were used to identify changes in a single species, six for changes in two species, and four for changes in three species (**Supplementary Table 4 contains the constraint matrices, Supplementary Document construction_of_constraint_matrices.pdf contains details on constructing the matrices**). Note that the contrast for changes in rhesus macaque will also identify changes that occurred in the human-chimpanzee-gorilla-orangutan internal branch. A value for each contrast *c* using constraint matrix *C* and variance-covariance matrix *Var*(*Cβ*) was calculated as *c* = [*Cβ*]*^T^*[*Var*(*Cβ*)]^−1^*Cβ*, where [*Var*(*Cβ*)]^−1^ denotes the matrix inverse of *Var*(*Cβ*). A p-value was calculated for each contrast using a chi-squared test to determine whether the accessibility for the species (or combination of species) of interest differed from rhesus macaque. We applied a Bonferroni correction for the 15 tests being conducted at each site. We took the contrast with the lowest, significant (p < 0.01), Bonferroni-adjusted p-value and the signs of the *β*values associated with the contrast to classify the pattern of differential accessibility across the species at the site. Note that while we chose the contrast with the lowest p-value, other contrasts may also be significant after multiplicity adjustment. For 2,454 (7% of differential sites) DHS sites, none of the p-values for the contrasts were less than 0.01, so the change was marked as “other”.

We have included supplementary files that contain the input data for the R script (**glm_analysis.input_file.txt**), the results from the analyses discussed in this paper (**glm_output_and_analsyis_results.txt**), and the field information for the input and output files (**input_and_output_file_field_information.xlsx**). Raw fastq files, bed files containing hg19 coordinates for the full read set, and bed files containing hg19 coordinates for the 20 million read subset are available under GEO accession GSE129034. The 20 million subset reads, DHS sites, and differential analysis classifications are available in a UCSC Genome Browser session at http://genome.ucsc.edu/s/ledsall/2019primate. All scripts for the data processing and analyses described above are available at http://github.com/ledsall/2019primate. Note that only the script named GO.step10.run_glm.R needs to be modified in order for researchers to use our method on their data sets. The GitHub repository contains a document (also included as **Supplementary Document modifying_the_GLM_analysis_R_script.pdf)** detailing the necessary modifications to that script. The other scripts in the repository are specific to the work reported here and are included for completeness and reproducibility.

### Classification of Differential Sites Using Multiple Pairwise Comparisons

As a separate analysis, we used edgeR (Robinson *et al*. 2010) (version 3.20.9) to perform multiple pairwise comparisons on the same input data used for the generalized linear model. We used these results as a comparison with our method. We calculated the normalization factors using the calcNormFactors command. We calculated the dispersion estimates using the estimateDisp command. We fit the model using the quasi-likelihood method (glmQLFit command with default parameters). We used the quasi-likelihood F-test (glmQLFTest command with default parameters) for four tests: human vs rhesus macaque; chimpanzee vs rhesus macaque; gorilla vs rhesus macaque; and orangutan vs rhesus macaque. We used the R function p.adjust to perform a Benjamini-Hochberg correction to adjust for multiple tests. A DHS site was considered differential if at least one species had a corrected p-value less than 0.01. A p-value threshold of 0.01 was selected to match the threshold used in the generalized linear model.

### Testing for Positive Selection and Determining Vertebrate Conservation

We performed selection analysis on both the differential and non differential DHS sites. We tested for selection on the species branches for human, chimpanzee, gorilla, and orangutan, and on the internal branches for human-chimpanzee and human-chimpanzee-gorilla.

As the stochasticity of the evolutionary process may be elevated in short alignments, we expanded each DHS site that was smaller than 300 bases up to 300 bases, while maintaining the size of any DHS site longer than 300 bases. We removed any sites that couldn’t be expanded due to gaps in the non-human genomes.

To investigate the extent of positive selection among the DHS sites, we used a branch-specific method we first developed in 2007 (Haygood *et al*. 2007) and recently improved (Berrio *et al*. 2019). Briefly, the method uses a likelihood ratio test based on the maximum likelihood estimates obtained from HyPhy (Pond *et al*. 2005). The branch of interest (e.g. human species branch) is used as the foreground and the rest of the tree is used as the background. The assumption for the background is the same for both the null and alternative models; specifically, neutral evolution and negative (purifying) selection are permitted but positive selection is not. In the null model, the assumption for the foreground is the same as the one for the background. In the alternative model, all three types of evolution are permitted (neutral evolution, negative selection, and positive selection) in the foreground. This method is highly sensitive and specific and can differentiate between positive selection and relaxation of constraint.

The method requires a 3kb reference alignment for each species that is used as a putatively neutral proxy for computing substitution rates. To generate this alignment, we first identified a set of functional regions on the human genome using annotations from the ENCODE project at UCSC (http://genome.ucsc.edu/encode/downloads.html) (ENCODE Project Consortium 2012) and annotations from the HoneyBadger2-intersect dataset from the ENCODE and Roadmap Epigenomics projects (https://personal.broadinstitute.org/meuleman/reg2map/HoneyBadger2-intersect_release/) (Roadmap Epigenomics Consortium *et al*. 2015). We used the set of 56,893 putative promoter regions; 1,598,323 putative enhancer regions; and 31,255 putative dyadic regions. We then masked the genomes using those functional regions, along with 5’ and 3’ UTRs, coding and non-coding RNAs, CpG repeats, microsatellite repeats, and simple repeats. Next, we extracted windows of 300 bases and excluded those with substitution rates that are too high or slow relative to the entire tree. Finally, we concatenated the set of these windows until we reached a length of 3kb (Berrio *et al*. 2019).

We used the PHAST library msa_split (Hubisz *et al*. 2011) to extract query regions from the UCSC Genome Browser Multiz Alignment of 100 Vertebrates (http://hgdownload.cse.ucsc.edu/goldenPath/hg19/multiz100way/maf/) (Blanchette *et al*. 2004) for the human, chimpanzee, gorilla, orangutan, and rhesus macaque genomes. For each DHS site (called a query site), we used HyPhy (Pond *et al*. 2005) to fit the null and alternative models and generate maximum likelihood values. We used a custom R script to compute the likelihood ratio, which was used as a test statistic for a chi-squared test with one degree of freedom to calculate a p-value. We classified a DHS site as under positive selection if the p-value was less than 0.05. We were unable to successfully run HyPhy on 12 sites due to unknown reasons and removed these regions from analysis.

We calculated the distribution of relative branch lengths for human, chimpanzee, gorilla, and orangutan (**Supplementary Figure 2**) for a random set of approximately 50,000 genomic regions. While the human, chimpanzee, and gorilla distributions are substantially similar, the orangutan distribution is much broader and shifted toward larger values. Whether this reflects a true biological difference or is an artifact of assembly quality or orthology assignment is not clear. In either case, this shift is sufficiently large that the substitution rate for orangutan biases the estimation of positive selection on that branch. Therefore, we excluded the orangutan branch from subsequent analysis. We also excluded the human-chimpanzee and human-chimpanzee-gorilla internal branches for two reasons. First, these internal nodes are predicted sequences rather than the observed sequences used for the external (species) branches and second, the short lengths of the internal branches often result in a divide-by-zero issue.

We then tested for significant enrichment of positive selection in different classes of DHS sites **(Supplementary Table 5)**. We performed Fisher’s exact tests using a test statistic of the number of DHS sites classified as under positive selection. We used a Bonferroni correction to adjust for the multiple tests performed.

To visualize the strength of selection, we computed the statistic *ζ* (zeta), representing the ratio of evolution, by calculating the ratio of the substitution rates in each query compared to the putatively neutral sites; we computed *ζ* for the human, chimpanzee, and gorilla species branches. This parameter is analogous to *ω* (omega), the ratio of dN/dS, where a value of *ω* < 1 indicates constraint or negative selection; a value of *ω* = 1 indicates neutrality; and a value of *ω* > 1 indicates positive selection.

We then tested whether the distributions of *ζ* differed between classes of differential DHS sites **(Supplementary Table 6)**. We performed Wilcoxon tests on human increased accessibility against the other classes of DHS sites with increased accessibility and used a Bonferroni correction to adjust for the multiple tests. Similarly, we performed Wilcoxon tests on human decreased accessibility against the other classes of DHS sites with decreased accessibility and used a Bonferroni correction to adjust for the multiple tests. Finally, we performed a Wilcoxon test on the non differential sites against the non functional sites defined above.

To determine the amount of vertebrate conservation, we computed the median value of the PhastCons scores for each DHS site using bedops (Neph *et al*. 2012), the UCSC 100-way PhastCons table (http://hgdownload.cse.ucsc.edu/goldenpath/hg19/phastCons100way) (Siepel *et al*. 2005; Pollard *et al*. 2010), and custom scripts. The PhastCons score represents the probability of a particular base being conserved. The values range from 0 to 1, with higher values representing an increased probability of conservation. Consistent with the original PhastCons paper (Siepel *et al*. 2005), we classified a DHS site as constrained if the median PhastCons score was above 0.9.

We then investigated whether the amount of conservation was similar between differential and non differential DHS sites. We used the percentage of constrained DHS sites as our test statistic and performed a Fisher’s exact test on non differential sites compared to three classes of differential DHS sites; specifically, 1) human accessibility changes; 2) chimpanzee accessibility changes; and 3) gorilla accessibility changes. We used a Bonferroni correction to adjust for the multiple tests.

### Intersection with Human Putative Regulatory Annotations

We characterized each DHS site as a proximal element, distal element, or unannotated region using the HoneyBadger2-intersect dataset from the ENCODE and Roadmap Epigenomics projects (https://personal.broadinstitute.org/meuleman/reg2map/HoneyBadger2-intersect_release/) (Roadmap Epigenomics Consortium *et al*. 2015). We used the putative promoter and enhancer regions as above, but did not use the putative dyadic regions. We used bedtools (Quinlan and Hall 2010) to identify DHS sites that overlapped the annotated promoters (which we characterized as “proximal elements”) and enhancers (which we characterized as “distal elements”). DHS sites that didn’t overlap promoters or enhancers were characterized as unannotated regions.

### Determining Cell Type Specificity

We characterized the cell type specificity of each DHS site by using bedtools (Quinlan and Hall 2010) to intersect it with DHS sites from 125 human cell types and tissues (http://hgdownload.cse.ucsc.edu/goldenpath/hg19/encodeDCC/wgEncodeAwgDnaseMasterSites/wgEncodeAwgDnaseMasterSites.bed.gz) (Thurman *et al*. 2012). We used a custom perl script (cluster_cell_types_in_bed_file.pl; available in the GitHub repository) and the ENCODE Cell Types metadata (https://genome.ucsc.edu/ENCODE/cellTypes.html) to remove cancerous cell lines and tissues (defined as having a value of “cancer” in the “Karyotype” column) and group the remaining samples into 32 tissue types (based on the value in the “Tissue” column). We assigned a score to each DHS site representing its cell type specificity. The score was calculated as 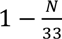, where *N* represents the number of cell types and tissues in which the DHS site is present (including the fibroblast cell line from this study). The score ranges from 0 for a DHS site present in all tissues and cell types to 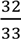 (which is approximately 0.97) for a DHS site present in only our dataset.

We then asked whether the distribution of cell type specificity scores varied between different classes of DHS sites **(Supplementary Table 7)**. We subset the DHS sites into those overlapping proximal elements and those overlapping distal elements. Within each subset, we performed Wilcoxon tests on the classes of DHS sites and used a Bonferroni correction to adjust for the multiple tests.

## Results

### Method Development to Identify and Classify Differential DNase I Hypersensitive Sites Across Multiple Primate Species

We developed a joint modelling method to allow us to quantitatively compare DNase I hypersensitive (DHS) sites across five primate species and identify a differential site with one statistical test. We then used contrasts to identify the species (or combination of species) with the most prominent change in accessibility compared to rhesus macaque (see **Materials and Methods**). The output from the model includes *β* values, which are analogous to log fold changes in conventional pairwise comparisons, and represent the difference in chromatin accessibility between a species and rhesus macaque (**Figure 1A-D, Supplementary Figures 3 and 4**).

**FIG. 1.**
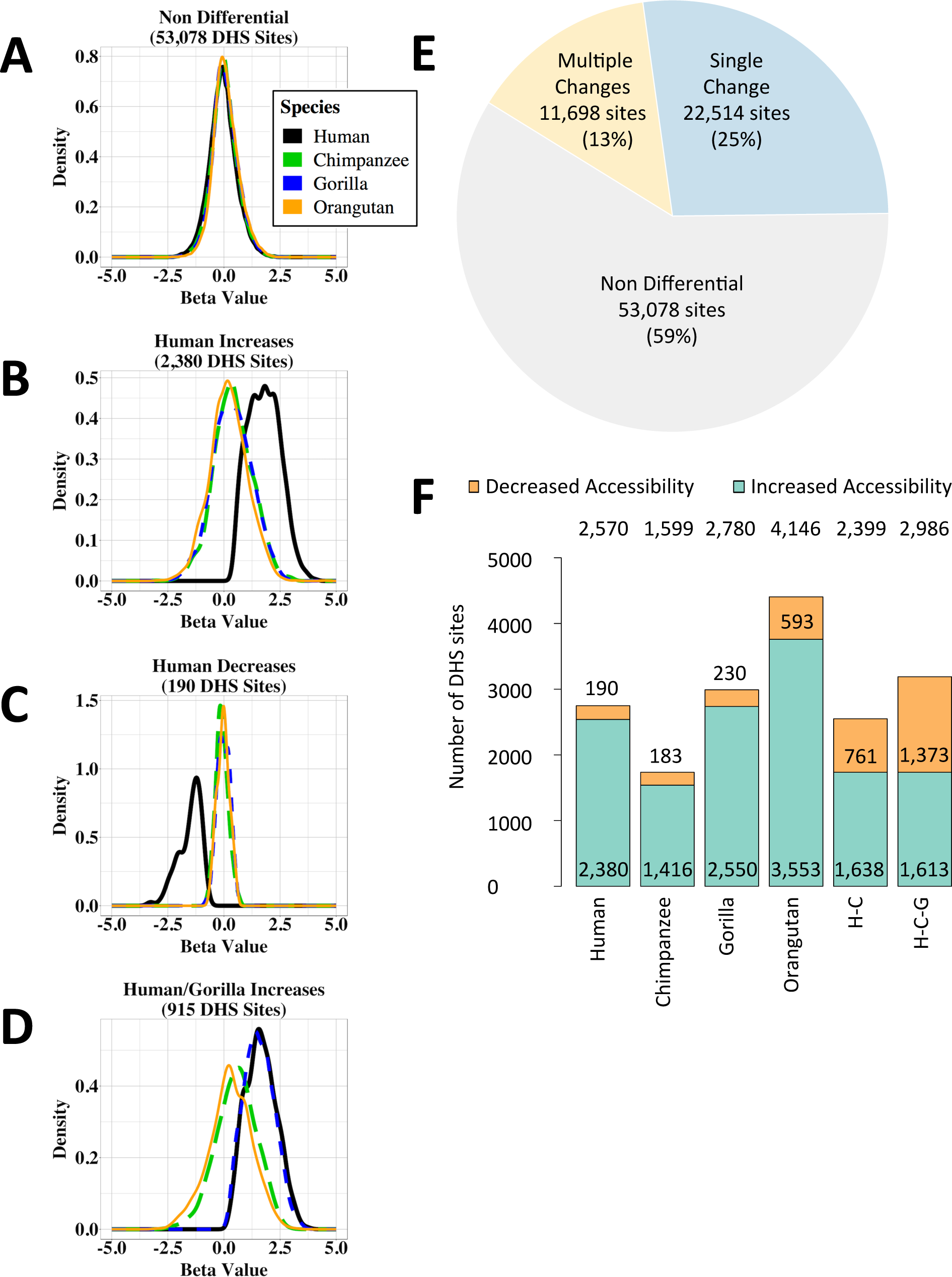
Classification of DHS sites. (A-D) Density plots showing the beta values for human (black), chimpanzee (green), gorilla (blue), and orangutan (orange). (A) Non differential sites. (B) Chromatin accessibility increases in human. (C) Chromatin accessibility decreases in human. (D) Chromatin accessibility increases in human and gorilla. (E) Pie chart showing the number and proportion of DHS sites that i) are non differential; ii) have accessibility changes likely due to a single event; and iii) have accessibility changes that are due to multiple events. Percentages are of the total number of DHS sites. Not shown: differential DHS sites that could not be classified due to insufficient power. (F) Bar chart showing the relative proportions of increases and decreases in accessibility. Numbers at the top are the total number of DHS sites in each category. Numbers in or just above the orange bar are the number of DHS sites with decreased accessibility. Numbers at the bottom of the green bar are the number of DHS sites with increased accessibility. H-C: Human-chimpanzee internal branch. H-C-G: Human-chimpanzee-gorilla internal branch.

**FIG. 2.**
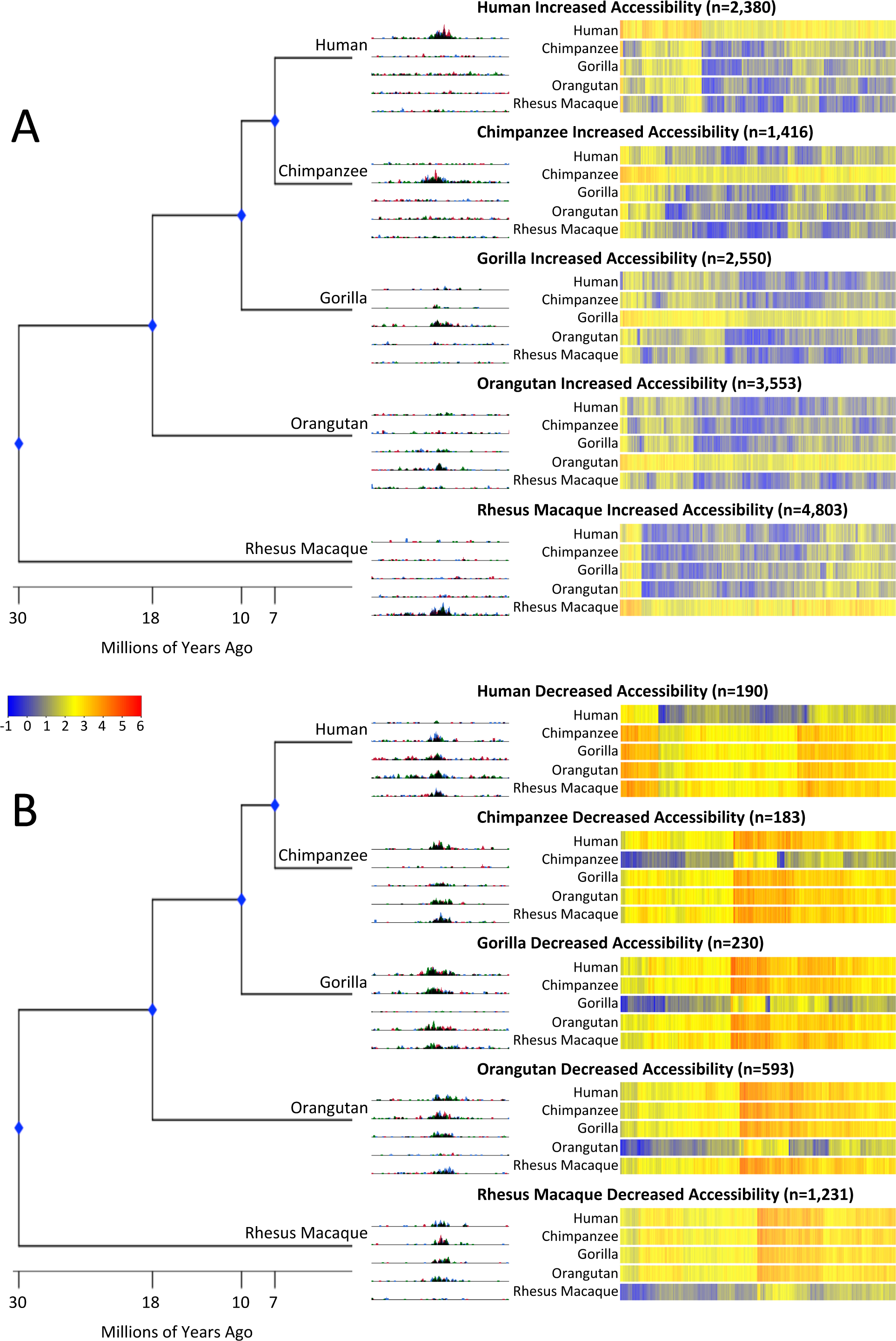
Chromatin accessibility changes in each species. Phylogenetic tree with divergence dates (to scale). UCSC Genome Browser screenshots of representative DHS sites for (A) increased accessibility and (B) decreased accessibility. Heatmaps of signal are rank-ordered DHS sites based on hierarchical clustering. Signal for the rhesus macaque species is equal to the rhesus macaque beta value. Signal for the non-rhesus macaque species is calculated by adding the rhesus macaque beta value to the species’ beta value.

**FIG. 3.**
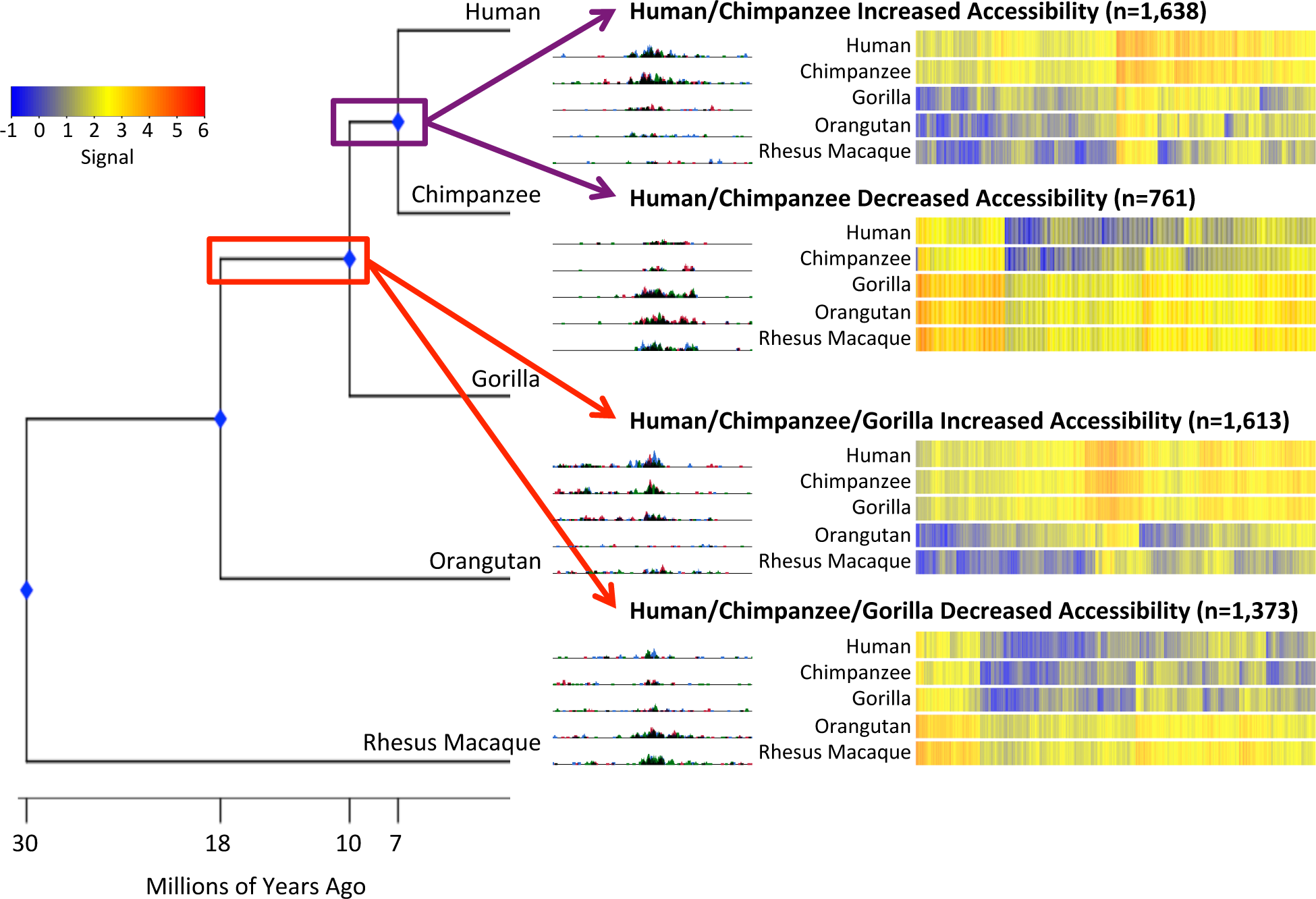
Internal branch changes in chromatin accessibility. Phylogenetic tree with divergence dates (to scale). UCSC Genome Browser screenshot of a representative DHS site changes that likely occurred before the human-chimpanzee split (top) and human-chimpanzee-gorilla split (bottom). Heatmaps of signal are rank-ordered DHS sites based on hierarchical clustering. Signal for the rhesus macaque species is equal to the rhesus macaque beta value. Signal for the non-rhesus macaque species is calculated by adding the rhesus macaque beta value to the species’ beta value.

**FIG. 4.**
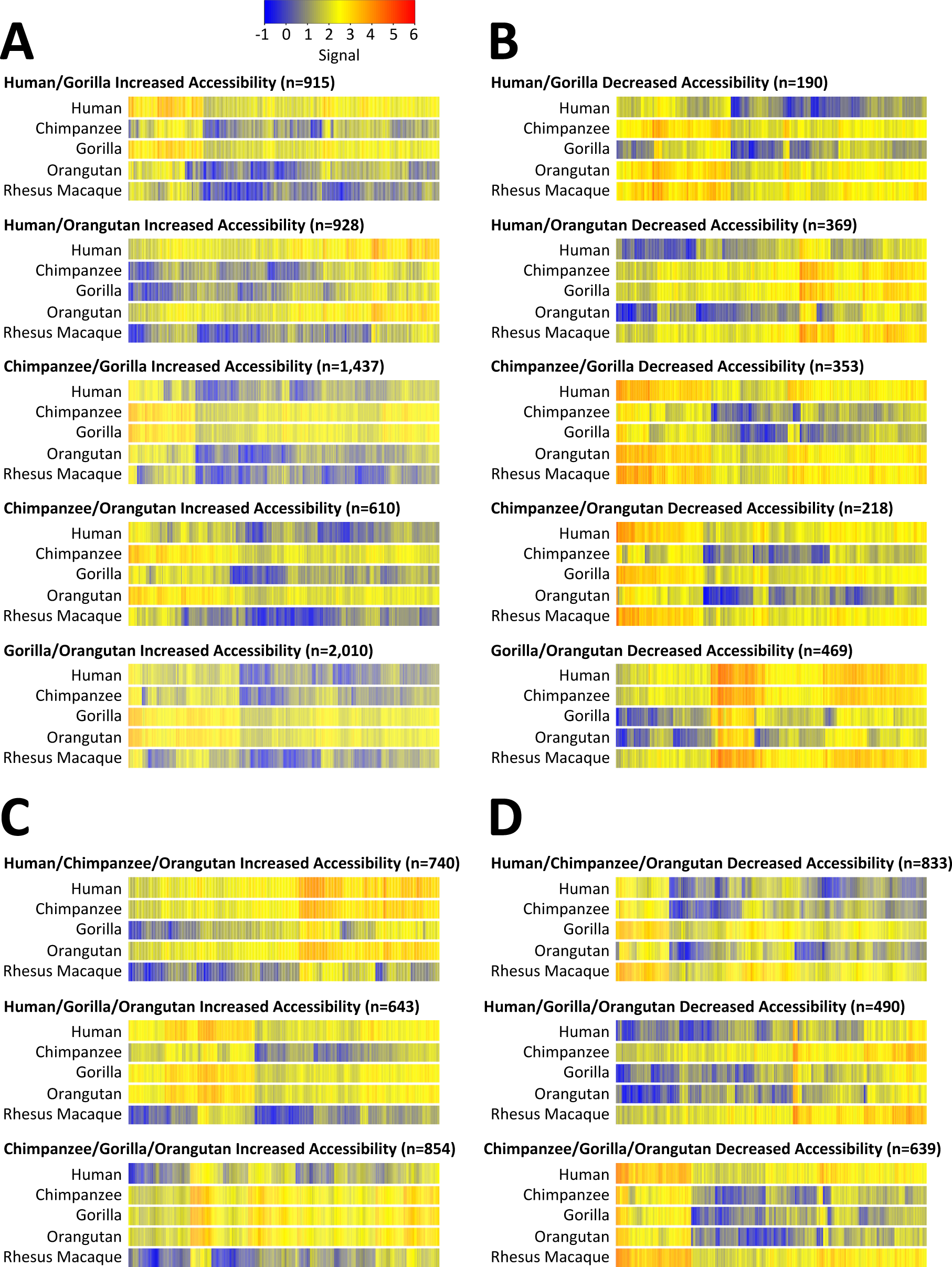
Changes in chromatin accessibility due to multiple events. Heatmaps of signal are rank-ordered DHS sites based on hierarchical clustering. Signal for the rhesus macaque species is equal to the rhesus macaque beta value. Signal for the non-rhesus macaque species is calculated by adding the rhesus macaque beta value to the species’ beta value. (A) Two species have increased chromatin accessibility relative to rhesus macaque. (B) Two species have decreased chromatin accessibility relative to rhesus macaque. (C) Three species have increased chromatin accessibility relative to rhesus macaque. (D) Three species have decreased chromatin accessibility relative to rhesus macaque.

Using this approach, we identified 89,744 total DHS sites that can be compared across all five species at 1:1:1:1:1 orthologous genomic regions (see **Materials and Methods**). As a first step in analyzing these data, we carried out a principal components analysis and found that the first principal component separated the single old world monkey (rhesus macaque) from the four great apes, while the second principal component recapitulated the phylogeny of the great apes (**Supplementary Figure 1**). Because we are drawing on the original data from Shibata *et al*. 2012 for three species (human, chimpanzee, rhesus macaque) and new data generated several years later for two species (gorilla, orangutan) (**Supplementary Table 1**), we also investigated whether batch effects would overwhelm the species signal by comparing principal components analyses performed with and without three additional chimpanzee samples generated more recently and reported in Pizzollo *et al*. 2018 (**Supplementary Table 1**). As shown in **Supplementary Figure 1**, the Pizzollo *et al*. chimpanzee samples cluster with the original Shibata *et al*. chimpanzee samples across the first four principal components (cumulative proportion of variance of 0.53), suggesting that biological signal is retained even when samples are prepared and sequenced years apart.

We performed additional analyses to determine the extent of technical and biological variation. For each species, we plotted the distribution of intra-species variation in normalized read counts across all of the DHS sites (**Supplementary Figure 5**). The distributions are highly similar across all five species, indicating that there are no major effects due to technical or biological differences. The similarity of the distributions also indicates a lack of systemic bias caused by differences in the quality and completeness of the genome assemblies. To further check the impact of biological differences, we compared the normalized read counts in all of the DHS sites between biological replicates within each species (**Supplementary Figure 6**). The biological replicates are highly concordant, even when replicates are from different sexes or ages.

Of the 89,744 total DHS sites, 53,078 (59%) are not statistically significantly different between species, 22,514 (25%) display a difference that likely resulted from a single chromatin accessibility change, and 11,698 (13%) display a difference due to multiple such changes **(Figure 1E**, **Table 1)**. For 2,454 differential sites, we were unable to determine the type of change, possibly due to low statistical power, and excluded them from subsequent analyses. Because we are using rhesus macaque as the outgroup, we are unable to differentiate between changes in the rhesus macaque species branch and changes in the human-chimpanzee-gorilla-orangutan internal branch.

**Table 1.** Number of DHS sites in each category

| **ALL DHS SITES** |  |  |
| --- | --- | --- |
|  | **Total** | **% of total** |
| Non differential (no changes) | 53,078 | 59% |
| Differential (one or more changes) | 36,666 | 41% |
| **TOTAL** | **89,744** |  |

| **SINGLE CHANGE AFFECTING SINGLE SPECIES** |  |  |  |  |
| --- | --- | --- | --- | --- |
|  | **Increases** | **Decreases** | **Total** | **% of differential** |
| Human | 2,380 | 190 | 2,570 | 7% |
| Chimpanzee | 1,416 | 183 | 1,599 | 4% |
| Gorilla | 2,550 | 230 | 2,780 | 8% |
| Orangutan | 3,553 | 593 | 4,146 | 11% |
| **TOTAL** | **9,899** | **1,196** | **11,095** | **30%** |

| **SINGLE CHANGE AFFECTING MULTIPLE SPECIES** |  |  |  |  |
| --- | --- | --- | --- | --- |
|  | **Increases** | **Decreases** | **Total** | **% of differential** |
| Human/chimpanzee | 1,638 | 761 | 2,399 | 7% |
| Human/chimpanzee/gorilla | 1,613 | 1,373 | 2,986 | 8% |
| Human/chimpanzee/gorilla/orangutan | 1,231 | 4,803 | 6,034 | 16% |
| **TOTAL** | **4,482** | **6,937** | **11,419** | **31%** |

| **MULTIPLE CHANGES AFFECTING TWO SPECIES** |  |  |  |  |
| --- | --- | --- | --- | --- |
|  | **Increases** | **Decreases** | **Total** | **% of differential** |
| Human/gorilla | 915 | 190 | 1,105 | 3% |
| Human/orangutan | 928 | 369 | 1,297 | 4% |
| Chimpanzee/gorilla | 1,437 | 353 | 1,790 | 5% |
| Chimpanzee/orangutan | 610 | 218 | 828 | 2% |
| Gorilla/orangutan | 2,010 | 469 | 2,479 | 7% |
| **TOTAL** | **5,900** | **1,599** | **7,499** | **20%** |

| **MULTIPLE CHANGES AFFECTING THREE SPECIES** |  |  |  |  |
| --- | --- | --- | --- | --- |
|  | **Increases** | **Decreases** | **Total** | **% of differential** |
| Human/chimpanzee/orangutan | 740 | 833 | 1,573 | 4% |
| Human/gorilla/orangutan | 643 | 490 | 1,133 | 3% |
| Chimpanzee/gorilla/orangutan | 854 | 639 | 1,493 | 4% |
| **TOTAL** | **2,237** | **1,962** | **4,199** | **11%** |

| **OTHER CHANGES** |  |  |
| --- | --- | --- |
|  | **Total** | **% of differential** |
| Not classified | 2,454 | 7% |

Consistent with our earlier study (Shibata *et al*. 2012), as well as studies by other groups (Reilly *et al*. 2015; Villar *et al*. 2015; Emera *et al*. 2016), the majority of the changes are increased accessibility rather than decreased accessibility (see **Discussion**). For changes on the species branches, there are approximately 10x the number of DHS sites with increased accessibility as DHS sites with decreased accessibility while changes on the internal branches have a ratio that is much less **(Figure 1F**, **Table 1)**.

### Changes in Chromatin Accessibility Detected in a Single Species

Using the methods described above, we identified 9,899 DHS sites with increased accessibility likely specific to a single species and 1,196 DHS sites with decreased accessibility likely specific to a single species **(Table 1)**. Heatmap overviews **(Figure 2)** of each class of increased accessibility and decreased accessibility show that these differences are not binary, but instead span the continuum from extremely large differences to those that represent more modest changes. Representative screenshots of individual genomic loci are shown in **Figure 2**.

Even though we can’t classify rhesus macaque-specific changes, we can identify sites where rhesus macaque is different from the other four species. We identified 4,803 sites that have increased accessibility in rhesus macaque relative to human, chimpanzee, gorilla, and orangutan **(Table 1)**. We identified 1,231 sites that have decreased accessibility in rhesus macaque compared to human, chimpanzee, gorilla, and orangutan **(Table 1)**.

### Changes in Chromatin Accessibility that Likely Occurred on Internal Branches

Our method allows us to identify ancient changes in chromatin accessibility that likely occurred as a single change on an internal branch. The contrasts we constructed and tested (see **Materials and Methods**) can identify changes that are present in human and chimpanzee (which likely occurred as a single change on the human-chimpanzee internal branch) and those that are present in human and chimpanzee and gorilla (which likely occurred as a single change on the human-chimpanzee-gorilla internal branch). We identified 1,638 DHS sites with increased accessibility and 761 DHS sites with decreased accessibility that likely occurred during the common lineage of human and chimpanzee **(Table 1)**. We identified 1,613 DHS sites with increased accessibility and 1,373 DHS sites with decreased accessibility that likely occurred before the split between human, chimpanzee, and gorilla **(Table 1)**. Heatmap overviews and representative screenshots of individual genomic loci are shown in **Figure 3**. As with accessibility changes in a single species, the heat map overviews show the changes are on a continuum rather than being binary.

### Multiple Changes in Chromatin Accessibility

In addition to detecting likely single changes in chromatin accessibility on either species or internal branches, we identified changes in chromatin accessibility that appear to have occurred multiple times, resulting in different combinations of chromatin accessibility patterns between species. There are many possible ways these differences could have happened and our method cannot determine if these changes resulted from multiple increases in accessibility, multiple decreases in accessibility, or a combination of increases and decreases (see **Discussion**).

We identified 5,900 DHS sites where two species display increased accessibility relative to rhesus macaque and 1,599 DHS sites where two species display decreased accessibility relative to rhesus macaque **(Table 1)**. We identified 2,237 DHS sites where three species displayed increased chromatin accessibility relative to rhesus macaque and 1,962 sites where three species displayed decreased accessibility relative to rhesus macaque **(Table 1)**. Heatmap overviews, showing a continuum of magnitudes of differences, and representative screenshots of individual genomic loci are shown in **Figure 4**.

### Comparison to Previous Study with Fewer Species

To test the accuracy of our new method for identifying differences in chromatin accessibility across five species, we compared our results with those from our previous study that used individual pairwise edgeR (Robinson *et al*. 2010) comparisons for human, chimpanzee, and rhesus macaque (Shibata *et al*. 2012). Using the species-specific calls from Shibata *et al*., we detected a high degree of concordance (**Supplementary Table 8**). Due to updates in the analysis pipeline, not all of the DHS sites that were previously characterized were also identified as DHS sites in this study **(Supplementary Table 9)**. The additional gorilla and orangutan DNase-seq data in this study allows us to fill in missing branch data and gauge the accuracy of our previous classification of human-specific or chimpanzee-specific changes. For 342 DHS sites that Shibata *et al*. characterized as human-specific increased accessibility, 245 (72%) are still characterized as human-specific increased accessibility even after including gorilla and orangutan, while 91 (27%) are now characterized as increased accessibility in human and at least one other species **(Supplementary Table 8)**. For 148 DHS sites that Shibata *et al*. characterized as human-specific decreased accessibility, 21 (14%) are still characterized as human-specific decreased accessibility even after including orangutan and gorilla, while 107 (72%) are now characterized as decreased accessibility in human and at least one other species **(Supplementary Table 8)**. A similar trend was detected for previously identified chimpanzee-specific changes **(Supplementary Table 8)**. For 1,154 DHS sites that Shibata *et al*. (2012) characterized as non differential between human, chimpanzee, and rhesus macaque, 5% (56) displayed changes in accessibility on the gorilla and/or orangutan branches that were not considered by Shibata *et al*., (**Supplementary Table 8**). Together, this indicates that adding chromatin accessibility data from additional primate species allows us to identify a substantial subset of DHS sites that have experienced changes in chromatin accessibility across multiple species during evolution.

### High Degree of Concordance with Conventional Multiple Pairwise Method

In order to compare our joint model to the conventional multiple pairwise method, we performed pairwise edgeR analyses using the same input data that was used for the generalized linear model (see **Materials and Methods**). We first checked whether the fold changes called by the joint model (represented on the natural log scale by the beta values) were consistent with the fold changes called by edgeR (**Supplementary Figure 7**). Overall, the joint model called more differential DHS sites (36,666) than the pairwise comparison (29,463); 26,093 DHS sites were called differential by both methods **(Table 2)**.

### DHS Sites with Decreased Accessibility are Enriched for Proximal Elements and DHS Sites with Increased Accessibility are Enriched for Distal Elements

After identifying and classifying DHS sites, we next determined their location in the human genome relative to previously annotated proximal and distal elements. We used the HoneyBadger2 annotations (see **Materials and Methods**), which are predicted promoters or enhancers based on histone marks identified in human cells and tissues as part of the Roadmap Epigenomics project (Roadmap Epigenomics Consortium *et al*. 2015). We overlapped these annotations to characterize each DHS site identified in this study as a proximal element, distal element, or unannotated region.

For DHS sites that are not differential between primate species, 22% (11,850) of these regions overlap proximal elements, 57% (30,371) overlap distal elements, and the remaining 20% (10,857) are unannotated **(Figure 5A**, **Supplementary Table 10)**. All DHS sites with increased accessibility relative to rhesus macaque display a substantially depleted amount of proximal element overlap compared to the non differential DHS sites (human: 3%; chimpanzee: 4%; gorilla: 9%; orangutan: 10%; human-chimpanzee: 3%; human-chimpanzee-gorilla: 5%) **(Figure 5A**, **Supplementary Table 10)**. Conversely, half of the classes of DHS sites with decreased accessibility relative to rhesus macaque overlap proximal elements to a similar degree as non differential DHS sites (human: 19%; chimpanzee: 22%; gorilla: 12%; orangutan: 21%; human-chimpanzee: 35%; human-chimpanzee-gorilla: 11%) **(Figure 5A**, **Supplementary Table 10**). These results indicate that decreased accessibility changes are more likely to be associated with proximal elements, while increased accessibility changes are more likely to be associated with distal elements. In every class of accessibility changes, there are substantially more distal than proximal elements, which is consistent with other studies (Schmidt *et al*. 2010; Villar *et al*. 2015).

**FIG. 5.**
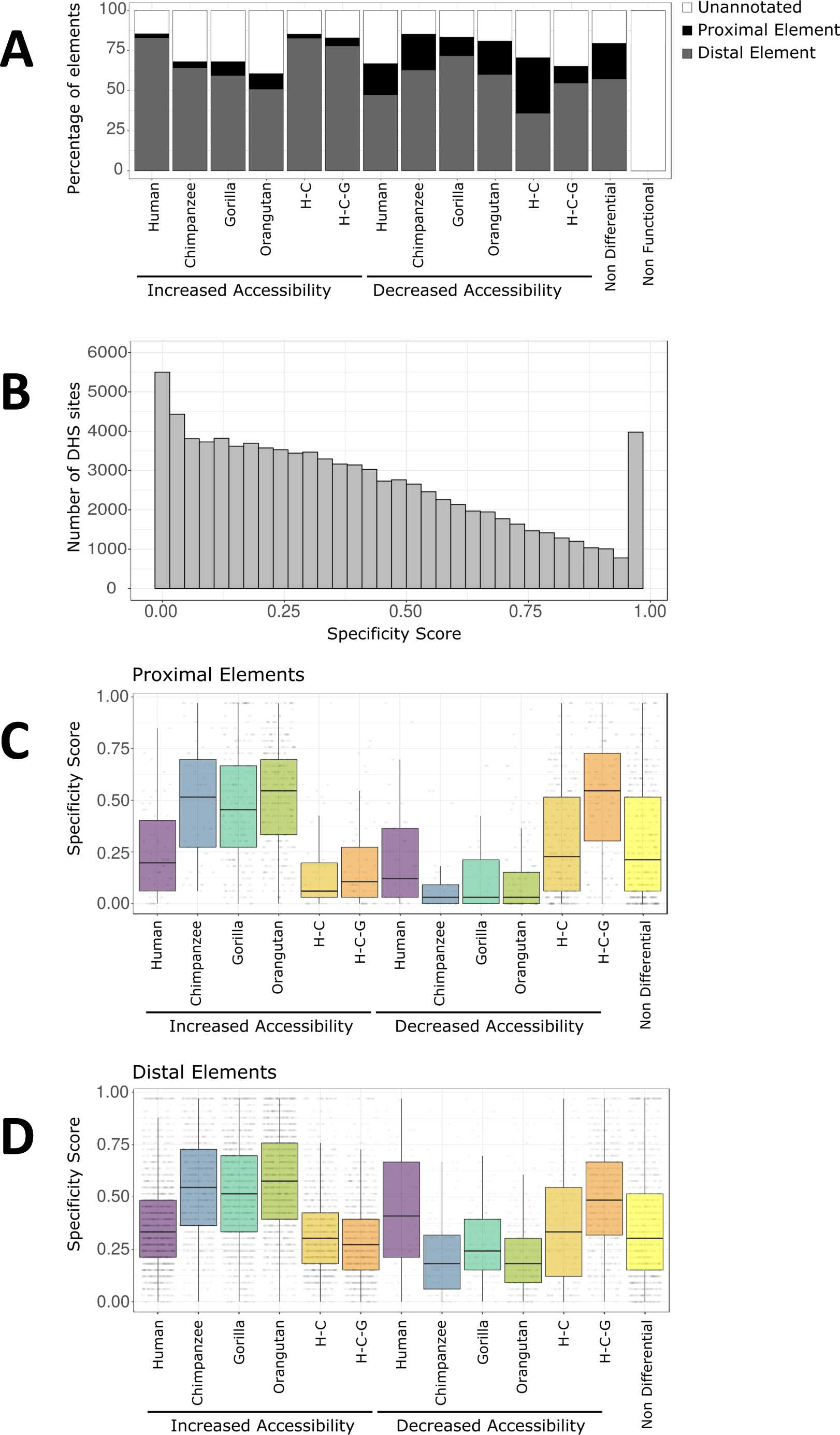
Chromatin accessibility changes relative to proximal/distal location and cell type specificity. (A) The percentage of proximal elements, distal elements, and unannotated elements for each category of DHS sites. (B) Histogram of specificity scores for DHS sites identified in this study compared to DHS sites detected in 32 different tissue and cell types (Thurman *et al*. 2012). A high specificity score indicates the DHS site is specific to a small number of cell types. A low specificity score indicates the DHS site is shared across many cell types. The DHS site categories are separated into proximal elements (C) and distal elements (D). H-C: human-chimpanzee internal branch. H-C-G: human-chimpanzee-gorilla internal branch.

We note that all of these proximal and distal annotations are from human tissues, which allows us to make specific inferences about comparisons only to human. There is not yet a similar Roadmap effort for non-human primate species. However, we find a high degree of overlap, which is likely due to non-human chromatin changes representing a continuum rather than being binary (e.g., open in non-human and completely closed in human). Classes of accessibility increases that include human have the lowest amount of overlap with unannotated regions of the genome. DHS sites with increased accessibility in chimpanzee, gorilla, or orangutan all have much higher overlaps with unannotated regions, with orangutan increased accessibility showing the highest degree of overlap with unannotated regions **(Figure 5A**, **Supplementary Table 10**). This is expected since orangutan is the most distantly related great ape species in our study. Similarly, we find that DHS sites with decreased accessibility in human have a higher overlap with unannotated regions compared to DHS sites with decreased accessibility in chimpanzee, gorilla, and orangutan. This is also expected since DHS sites with decreased accessibility in non-human primates will by definition have higher chromatin accessibility signals in human fibroblasts.

### Evolutionary Changes in Accessibility are Associated with Cell Type Specificity

We calculated cell type specificity (see **Materials and Methods**) for the DHS sites by comparing them to a much larger set of DHS sites detected in 32 different human cell and tissue types (Thurman *et al*. 2012). A cell type specificity score close to 1 indicates the DHS site is present in only a few of the 32 tissues and cell types, while a score near 0 indicates that the DHS site is present in almost all of the 32 tissues and cell types.

As with the proximal and distal annotations, we can make inferences about evolutionary changes in chromatin accessibility only for DHS sites that overlap the human annotations. The union set of the DHS sites we identified show a continuum of cell type specificity scores with DHS sites from different human cell types (**Figure 5B**). 5,502 (6%) of our DHS sites overlapped DHS sites found in all 32 tissues and cell types and 3,976 (4%) of our DHS sites were not found in any of the previously tested tissues and cell types.

We then analyzed the distribution of cell type specificity scores in distal and proximal DHS sites that displayed changes in chromatin accessibility. In general, distal elements have higher specificity scores than proximal elements **(Figure 5C and 5D)**, consistent with previous studies (Thurman *et al*. 2012).

For proximal elements showing increases in accessibility, tissue specificity is higher on all four species branches than on the two internal branches (one sided Wilcoxon test comparing pooled distributions of external vs internal; *P* = 1.64×10^−31^) **(Figure 5C)**. The opposite pattern is evident for decreases in accessibility (one sided Wilcoxon test comparing pooled distributions of external vs internal; *P* = 8.52×10^−30^) **(Figure 5C)**. Since changes on the internal branches are more ancient than those on external branches, this result hints at the possibility that degree of chromatin accessibility is positively correlated with broader utilization across cell types. One possible explanation is that increases in chromatin accessibility raise the likelihood that a proximal regulatory element is co-opted for use by another tissue. The same trends are observed for distal elements which have higher tissue specificity scores than proximal elements (**Figure 5D**). This is expected since distal chromatin accessible sites are more likely to be cell type specific than proximal elements (Xi *et al*. 2007; Thurman *et al*. 2012).

For proximal elements showing changes in chromatin accessibility, the human branch shows lower cell type specificity compared to the three other species for accessibility increases (one sided Wilcoxon test with Bonferroni correction; *P_H:C_* = 8.77×10^−7^; *P_H:G_* = 7.34×10^−9^; *P_H:O_* = 7.17×10^−14^) and higher cell type specificity for accessibility decreases compared to chimpanzee and orangutan (one sided Wilcoxon test with Bonferroni correction; *P_H:C_* = 0.008; *P_H:G_* = 0.19; *P_H:O_* = 0.011) **(Figure 5C)**. A similar pattern is present for distal elements, for both increases and decreases in accessibility (one sided Wilcoxon test with Bonferroni correction; Increases: *P_H:C_* = 3.27×10^−75^; *P_H:G_* = 1.42×10^−64^; *P_H:O_* = 1.10×10^−147^; Decreases: *P_H:C_* = 1.61×10^−7^; *P_H:G_* = 3.56×10^−4^; *P_H:O_* = 1.38×10^−10^) **(Figure 5D)**. This may reflect an ascertainment bias arising from relying on human tissue comparisons for the cell type specificity score.

### Selection Within DHS Sites Showing Chromatin Changes

To investigate the evolutionary significance of species-specific changes in chromatin accessibility, we tested each DHS site for signatures of positive selection on the human, chimpanzee, and gorilla branches separately (see **Materials and Methods**). Testing for positive selection required additional filtering of DHS sites (see **Materials and Methods**), resulting in a reduced set of 87,431 DHS sites used in this analysis. The figure of merit in these analyses is ζ (zeta), the ratio of substitution rates within a DHS site on a given branch compared to the substitution rates for a collection of proxy neutral sites (Wong and Nielsen 2004; Haygood et al. 2007; Haygood et al. 2010). Similar to the analogous and more familiar ω (omega), high values of ζ indicate positive selection, values near 1 indicate neutrality, and low values indicate negative selection.

Putative non functional elements display a relatively tight distribution of ζ on the human branch centered around 1 (**Figure 6A**), confirming they are a good proxy for neutral evolution in non-coding regions of the genome. Non differential DHS sites have a distribution of ζ on the human branch that is centered significantly below 1 (one sided Wilcoxon test; P = 1.57×10^−283^) (**Figure 6A**), consistent with ongoing negative selection. Additionally, the distribution of ζ values is much broader for non differential DHS sites compared to putative non functional sites, with a small fraction showing elevated substitution rates on the human branch that are consistent with positive selection.

**FIG. 6.**
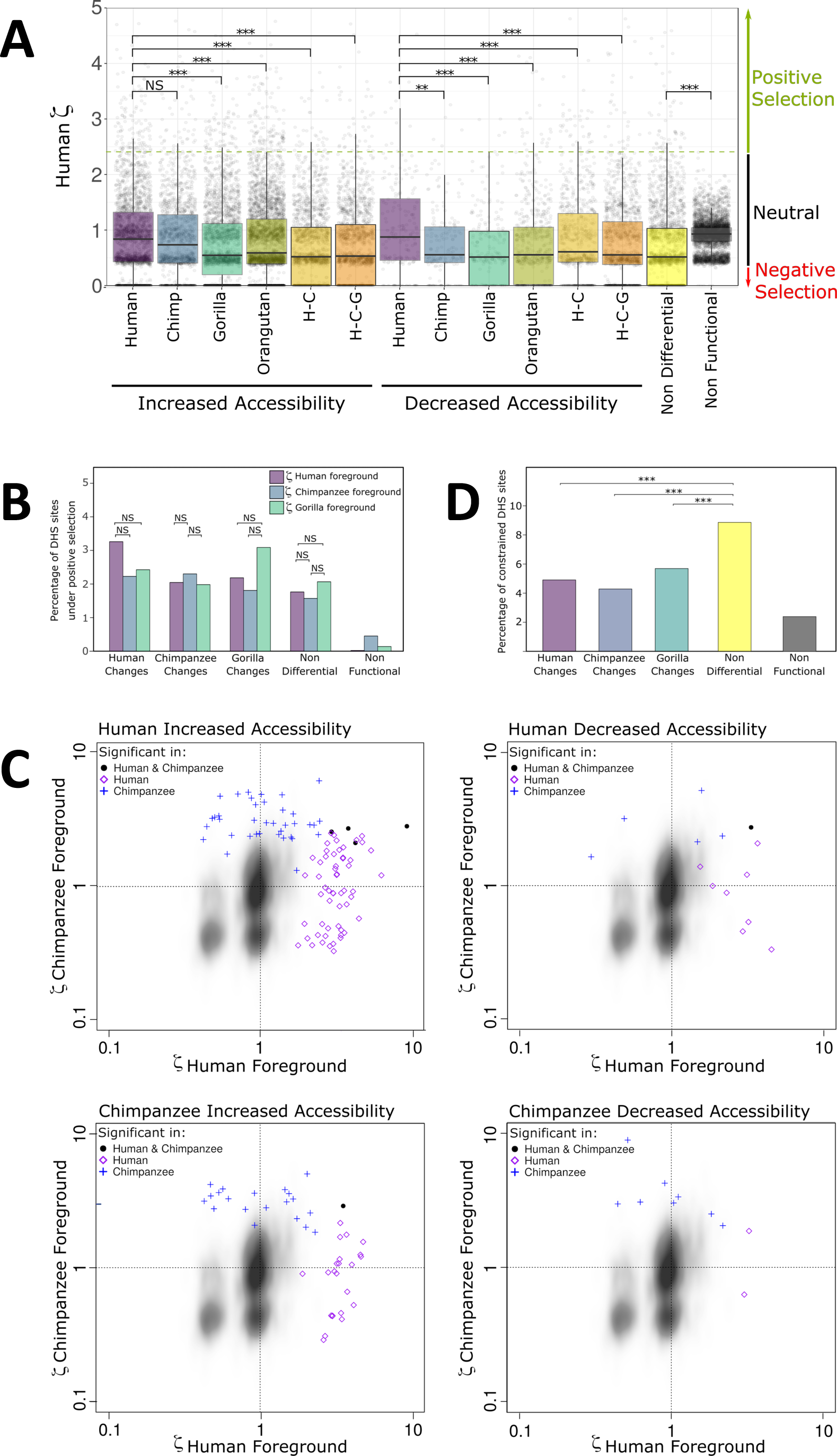
Effect of natural selection on increases and decreases in chromatin accessibility. (A) Distribution of the ratio of evolution ζ (zeta) in the human branch for DHS sites. The dashed green line depicts the critical value where the human zeta value becomes significant (p < 0.05). Zeta values around 1 are expected to be neutral and below 1 are expected to be constrained. (B) Percentages of DHS sites that are significant for positive selection (p < 0.05). Each DHS site was tested with 3 different foregrounds: human, chimpanzee, and gorilla. (C) Scatterplots of zeta values for DHS sites with significant positive selection on the human branch (purple diamond), the chimpanzee branch (blue cross), or both the human and chimpanzee branches (black solid circle). Zeta values for the human branch are on the x-axis and zeta values for the chimpanzee branch are on the y-axis. The kernel density depicts non functional sites. (Top left) increased accessibility in human. (Top right) decreased accessibility in human. (Bottom left) increased accessibility in chimpanzee. (Bottom right) decreased accessibility in chimpanzee. (D) Percentages of DHS sites that are highly constrained (median vertebrate PhastCons > 0.9). H-C: human-chimpanzee internal branch. H-C-G: human-chimpanzee-gorilla internal branch. ***: p-value < 0.001. **: p-value < 0.01. *: p-value < 0.05. #: p-value < 0.1.

DHS sites with a change in chromatin accessibility on the human branch have positively-shifted distributions of ζ on the human branch relative to non differential DHS sites (**Figure 6A**). This suggests that both increases and decreases in accessibility are accompanied by enrichment for a combination of relaxed selection and positive selection on the same branch. As expected, this enrichment on the human branch is less pronounced when the accessibility change occurs on a different branch of the phylogeny: the distributions of ζ on the human branch are higher when the chromatin accessibility change occurred on the human branch rather than the gorilla or orangutan branches, and this is true for both increases and decreases (one sided Wilcoxon test with Bonferroni correction; Increases: P_H:C_ = 0.12, P_H:G_ = 4.74×10^−11^, P_H:O_ = 1.99×10^−6^, P_H:H-C_ = 9.13×10^−14^, P_H:H-C-G_ = 3.89×10^−11^; Decreases: P_H:C_ = 4.52×10^−3^, P_H:G_ = 9.25×10^−5^, P_H:O_ = 1.15×10^−4^, P_H:H-C_ = 0.03, P_H:H-C-G_ = 2.62×10^−4^), although these differences are all modest in magnitude (**Figure 6A**).

For the human accessibility changes, we tested for enrichment of positive selection on the human branch relative to the chimpanzee and gorilla branches. We performed a similar comparison for the chimpanzee accessibility changes by testing for enrichment of positive selection on the chimpanzee branch relative to the human and gorilla branches. Finally, for the gorilla accessibility changes, we tested for enrichment of positive selection on the gorilla branch relative to the human and chimpanzee branches (**Figure 6B**). Although none of the Fisher’s exact tests were significant after Bonferroni correction for the total number of foreground branches considered (n=6), the data trends in the expected patterns (e.g., human changes have more selection on the human branch). As a control, non differential DHS sites show no significant differences in positive selection between branches (**Figure 6B**). Additionally, putative non functional sites in the genome do not display an enrichment of positive selection (**Figure 6B**). These results suggest that evolutionary changes in chromatin accessibility between species are phylogenetically correlated with an enrichment of positive selection.

Next, we investigated the converse: whether signatures of positive selection on individual regulatory elements are generally limited to branches where the change in accessibility occurred. This is clearly not the case: DHS sites with increased accessibility on the human branch show positive selection on the chimpanzee branch almost as often as on the human branch (**Figure 6C**). The same pattern is evident for increased accessibility on the chimpanzee branch and for decreased accessibility on the human or chimpanzee branch (**Figure 6C**). These results suggest that positive selection may act on the DNA sequence of DHS sites in ways that do not affect chromatin accessibility. Interestingly, some DHS sites show evidence of positive selection on both branches (**Figure 6C**).

On average, 5% of DHS sites that show accessibility changes in human, chimpanzee, or gorilla are highly constrained in sequence evolution, with nucleotide substitution rates that are significantly lower than the neutral expectation across vertebrates (**Figure 6D**). In contrast, DHS sites that are not differential are enriched for highly constrained sites in comparison to accessibility changes in human, chimpanzee, or gorilla (Fisher’s exact test, one-sided with a Bonferroni correction for 3 comparisons; P_ND:H_ = 1.23×10^−9^; P_ND:C_=1.47×10^−8^; P_ND:G_=5.10×10^−6^) (**Figure 6D**).

Together, these results suggest that positive selection contributes to chromatin accessibility increases and decreases, while purifying selection contributes to the conservation of non differential DHS sites.

## Discussion

We developed a new method that uses a negative binomial generalized linear model to identify regions of differential chromatin accessibility across multiple species. This method does not rely on thresholding and is therefore able to detect subtle differences in degree of chromatin accessibility that are obscured using conventional approaches. In addition, our method jointly models the data across all species and replication. We carry out a single global test for any difference among species at a particular genomic location that acts as a “gatekeeper” (Dmitrienko and Tamhane 2007). In contrast, the conventional approach of multiple pairwise comparisons requires correcting for the number of pairwise comparisons, which scales exponentially and thus significantly decreases sensitivity. For example, in this study, the joint model method required 89,744 tests while the conventional method required 358,976 tests. As shown in Table 2, the joint model identified 7,203 (8% of all DHS sites) more differences among species than the conventional pairwise approach. This is due in part to a lower multiple comparisons burden, which in turn allows the method to detect more subtle quantitative changes.

**Table 2.** Comparison of joint model classifications with pairwise classifications NOTE. —(a) Percentages are of the total number of pairwise classifications. (b) Percentages are of the total number of joint model classifications.

| **ALL DHS SITES** |  |  |  |  |  |  |
| --- | --- | --- | --- | --- | --- | --- |
|  | **Either**<br>**Method** | **Both**<br>**Methods** | **Pairwise**<br>**Only (a)** | **Joint Model**<br>**Only (b)** | **Pairwise**<br>**Total** | **Joint**<br>**Model**<br>**Total** |
| Non Differential | 63,651 | 49,708 | 10,573 18% | 3,370 6% | 60,281 | 53,078 |
| Differential | 40,036 | 26,093 | 3,370 11% | 10,573 29% | 29,463 | 36,666 |

A joint model also provides a more principled approach to dealing with cases where multiple state changes have evolved among species. This is not a problem when only two ingroup species are analyzed, but as the number of ingroup species rises it becomes increasingly more difficult to reconstruct the history of state changes across the phylogeny. In addition, the number of state changes within any given locus will on average rise as the number of taxa analyzed rises. With pairwise comparisons, the reconstruction of state changes is based on *post hoc* review of several independent comparisons, each of which only consider data from two species. Our approach draws on all the available data, providing a more principled approach to identification of state changes. It does not, by itself, reconstruct the most likely history of state changes across the phylogeny, but it does estimate a figure of merit (beta values) that can be input into conventional tools for character state reconstruction conditional on a phylogeny.

Our approach tests for quantitative differences among species and incorporates phylogenetic topology after chromatin accessibility changes have been identified. As a result, it is complementary to conventional methods for inferring inter-species changes in quantitative traits. It is also possible to use our approach when phylogeny is ambiguous.

Here we applied the joint model to DNase-seq data from cultured skin fibroblasts from five primate species. While the majority of DHS sites (59%) were not quantitatively distinct between species, we identified 36,666 DHS sites with significant differences in chromatin accessibility between human, chimpanzee, gorilla, orangutan, and rhesus macaque. Of those, 61% are likely the result of a single change in chromatin accessibility that occured on either an internal or external branch, while the remainder imply multiple changes in accessibility.

Our results show a high degree of overlap with a conventional analysis using pairwise comparisons and include modest changes that the conventional method was not able to detect. Our results are also largely congruent with our earlier study (Shibata *et al*. 2012) that used a threshold-based multiple pairwise comparison approach and considered three primate species (human, chimpanzee, and rhesus macaque). Here, the use of five species provides additional confidence in the identification of species-specific accessibility changes and also allows for the identification of accessibility changes that likely occurred multiple times throughout evolution. For these multiple changes, the method we developed does not characterize how exactly they occurred. That characterization will be the subject of future work using a likelihood analysis that incorporates the phylogenetic information and models evolutionary processes (Felsenstein 1973; Hansen 1997; Felsenstein 2008; Paradis and Schliep 2019).

As mentioned in the results, we identified substantially more accessibility increases than decreases. It seems in principle unlikely that increases and decreases in accessibility actually occur at such different rates, since, if true, primate genomes would eventually become saturated with open chromatin regions. The same asymmetry was observed previously by us (Shibata *et al*. 2012) and other groups (Villar *et al*. 2015; Reilly *et al*. 2015; Emera *et al*. 2016) using conventional pairwise comparisons and thresholding, so the source is unlikely to lie in the method we developed and describe here. Instead, it seems likely that the asymmetry is an ascertainment bias that derives, in part, from unequal statistical power to call increases and decreases, though the exact basis of the bias is currently unclear.

Finding so many DHS site differences in non-human primates is a fascinating result with implications for understanding the evolution of transcriptional regulation. Nevertheless, we also suggest that the results describing cell type specificity should be interpreted carefully. One non-biological possible scenario for such enrichment is an ascertainment bias in our analyses due to the cell type specificity score being based entirely on data from human, a limitation imposed by the current lack of comparable cell type specificity data from other primate species. Although the patterns of positive selection that we detected are consistent with expectations, none of the tests found statistically significant enrichment on the human, chimpanzee, or gorilla branches after correcting for multiple testing. This may be due to our method of positive selection detection relying on human functional annotations to identify proxy neutral regions, which may result in a loss of power with increasing phylogenetic distance.

Interestingly, our results suggest that DHS sites are not homogenous from either a functional or an evolutionary perspective. Those near transcription start sites (including likely core promoter regions) differ from DHS sites that are distant (including classic enhancers and other kinds of distal elements) in several regards. Compared with proximal DHS sites, gains in chromatin accessibility in distal sites are more likely to show signatures of positive selection on the same branch, as might be expected if these DHS sites are contributing to changes in gene regulation. DHS sites that are not differential between the species surveyed are enriched in conserved nucleotides, consistent with greater functional constraint. These and other trends we observed suggest that functional constraints and opportunities differ markedly among classes of DHS sites. Additional studies will be needed to delineate these distinct classes of likely regulatory elements and to understand how evolutionary mechanisms operate on their chromatin accessibility and underlying DNA sequence.

Functional characterization studies will be necessary to understand these regions and their contribution to gene expression patterns and organismal traits. High-throughput reporter assays such as MPRA (Klein *et al*. 2018) and population STARR-seq (Vockley *et al*. 2015) can quantify the impact of these differentially utilized regulatory regions, as well as variants within these regions. In addition, methods such as CRISPR (Diao *et al*. 2017; Klann *et al*. 2017) can characterize the impact of these regions in their natural context, including identifying the correct target gene(s) for these regulatory elements. Finally, additional replicates from these species can provide characterization of biological variability within each species. While obtaining data from additional tissues for primate species is not possible for most tissues, generation of induced pluripotent stem cells (iPSCs) followed by differentiation (Gallego Romero *et al*. 2018), will provide insights into how these differential chromatin signals translate into different cell types across many species.

While we used our joint model method to identify and classify differences in chromatin accessibility between five species, we believe this strategy can be used for quantitative comparisons across tissues, cell types, time-series, and similar experiments. In addition to DNase-seq, we expect this method can be readily applied to any count-based data type such as RNA-seq, ATAC-seq, and ChIP-seq because the input is a table of read counts. The procedure to generate this input table will vary between the different types of assays, but once the input table is generated, the procedure is the same regardless of the source of the data. The identification of differential sites using this method is also easily adaptable to more than five groups, as it only requires changing the design matrices.

## Acknowledgments

We would like to thank Terry Gaasterland for her help developing the tiered mapping approach. We thank the Duke Sequencing and Genomic Technologies Shared Resource for sequencing. This work was supported by the National Science Foundation [HOMINID 0827552 to G.A.W]; the National Institute of Mental Health at the National Institutes of Health [5R01MH105472 to G.E.C and G.A.W]; and a generous donation from Dr. Howard Clark. This work used a high-performance computing facility partially supported by grant 2016-IDG-1013 (“HARDAC+: Reproducible HPC for Next-generation Genomics”) from the North Carolina Biotechnology Center.

