## Supplementary material for "Evaluating chromatin accessibility differences across multiple primate species using a joint modelling approach": construction_of_constraint_matrices.pdf

The contrasts compare the geometric means of the non-macaque species to each other. To determine the coefficients of the constraint matrices, it is helpful to work out the comparisons using equations containing the geometric means. A few things to note:

- Geometric means use multiplication instead of addition.
- The terms on each side of the equation are raised to the power of $1/n$ where $n$ is the number of terms on that side of the equation.
- The coefficients on the left hand side of the equation are entered into the constraint matrix as positive numbers.
- The coefficients on the right hand side of the equation are entered into the constraint matrix as negative numbers.
- Our constraint matrices are in the order: (1) human; (2) chimpanzee; (3) gorilla; (4) orangutan

Example: Human-specific changes. The constraint matrix is [1 -¼ -¼ -¼]

$$\mu_H \neq (\mu_C \mu_G \mu_O \mu_M)^{1/4}$$

$$e^{\beta_M + \beta_H} \neq (e^{\beta_M + \beta_C}\ e^{\beta_M + \beta_G}\ e^{\beta_M + \beta_O}\ e^{\beta_M})^{1/4}$$

$$e^{\beta_M} e^{\beta_H} \neq e^{\beta_M}(e^{\beta_C}\ e^{\beta_G}\ e^{\beta_O})^{1/4}$$

$$e^{\beta_H} \neq (e^{\beta_C}\ e^{\beta_G}\ e^{\beta_O})^{1/4}$$

$$log(e^{\beta_H}) \neq log((e^{\beta_C}\ e^{\beta_G}\ e^{\beta_O})^{1/4})$$

$$\beta_H \neq \frac{\beta_C + \beta_G + \beta_O}{4}$$

Example: Human/Chimpanzee changes. The constraint matrix is [½ ½ -⅓ -⅓]

$$(\mu_H \mu_C)^{1/2} \neq (\mu_G \mu_O \mu_M)^{1/3}$$

$$(e^{\beta_M + \beta_H}\ e^{\beta_M + \beta_C})^{1/2} \neq (e^{\beta_M + \beta_G}\ e^{\beta_M + \beta_O}\ e^{\beta_M})^{1/3}$$

$$e^{\beta_M}(e^{\beta_H}\ e^{\beta_C})^{1/2} \neq e^{\beta_M}(\ e^{\beta_G}\ e^{\beta_O})^{1/3}$$

$$(e^{\beta_H}\ e^{\beta_C})^{1/2} \neq (\ e^{\beta_G}\ e^{\beta_O})^{1/3}$$

$$log((e^{\beta_H}\ e^{\beta_C})^{1/2}) \neq log((\ e^{\beta_G}\ e^{\beta_O})^{1/3})$$

$$\frac{\beta_H + \beta_C}{2} \neq \frac{\beta_G + \beta_O}{3}$$
