## Supplementary material for "Evaluating chromatin accessibility differences across multiple primate species using a joint modelling approach": modifying_the_GLM_analysis_R_script.pdf

### Modifying the GLM Analysis R Script (GO.step10.run_glm.R)

#### Overview

This script was designed to analyze DNase-seq data from 3 biological replicates in each of 5 species (human, chimpanzee, gorilla, orangutan, and macaque). We used macaque as a baseline and all of the results are relative to it (e.g. human read counts are increased relative to macaque). In linear model terminology, macaque is referred to as the intercept.

It should be possible to run this script on any count-based data set, such as those generated by ChIP-seq, ATAC-seq, and RNA-seq. You will need to modify the script to analyze your data. Details on the necessary modifications are included in this document. Modifications are necessary for different numbers of replicates, different numbers of groups, and group names. Modifications are not necessary when the only thing changing is the type of experiment (e.g., ChIP-seq vs. RNA-seq).

The script is provided under CC-BY. If used in a publication, either as-is or modified, please cite Edsall et al., 2019.

#### Required libraries

The script requires 2 libraries: DSS and MASS.

- DSS
  - “Dispersion shrinkage for sequencing data”
  - It contains functions to calculate the dispersion and normalization offset parameters.
  - It is available from Bioconductor.
  - Citation: Wu H, Wang C, Wu Z. 2013. A new shrinkage estimator for dispersion improves differential expression detection in RNA-seq data. Biostatistics 14(2):232-243.
- MASS
  - It is required for the negative.binomial function.
  - It is available from CRAN.

#### Analysis is performed in three main parts

The first part uses DSS to calculate the dispersion and normalization offset parameters. These parameters are used in the glm function calls in the next two parts. All of the sites are analyzed with one command.

The second part identifies the differential sites and updates a “gateway flag”. This is a character flag that uses `GNS` for non significantly differential sites and `GD` for differential sites. A for loop iterates through the sites. Each iteration calls the `glm` function and calculates a raw p-value. After the loop finishes, a Benjamini-Hochberg correction is performed using the `p.adjust` function.

The third part classifies each differential site. It uses a for loop to iterate through all of the sites and skips ones that are non differential. For the differential sites, the loop calls the `glm` function and extracts the variance-covariance matrix and performs constraint tests.

#### Script layout

The script has 10 sections. Before each section, there is a comment block with the name and a brief overview. The name always starts with “SECTION” and you can quickly move to the next section by searching for the word “SECTION”.

###### SECTION 1 OF 10: SETUP

This section is for loading libraries, setting global options and variables, and opening a connection to a log file. The threshold for p-value significance is set here; we use a value of .01.

Modifications

- The threshold for p-value significance
- The name of the log file

###### SECTION 2 OF 10: READ THE SCORE FILE AND PREPARE IT FOR THE GLM FUNCTION

The input file needs to contain non-normalized integer scores. We recommend including labels for each site, such as chromosomal coordinates or transcript identifiers. Our file has the samples in the columns and the locations in the rows, but the script can be easily modified for an input file with the samples in the rows and the locations in the columns.

Details of our input file

- The samples in the columns and the locations are in the rows.
- The columns are tab-separated.
- There isn’t a header.
- There are 19 columns.
- Columns 1-3 contain location information (chromosome, start, end).
- Columns 4-6 contain scores for the human samples.
- Columns 7-9 contain scores for the chimpanzee samples.
- Columns 10-12 contain scores for the gorilla samples
- Columns 13-15 contain scores for the orangutan samples.
- Columns 16-18 contain scores for the macaque samples.
- Column 19 contains a flag indicating overlap with regions from a previous study.

Modifications

- The name of the file
- The `read.table` command if there is a header or the columns are not tab-separated
- The column names
- The row names
- The transpose command
- The row names of the transposed matrix

**SECTION 3 OF 10: RUN DSS TO GET THE DISPERSION AND NORMALIZATION PARAMETERS**

The DSS package requires the specification of a design matrix. The matrix associates a group name to each replicate. In our case, we used the species names as the group names. The group names are completely arbitrary. Each replicate needs an entry in the design matrix.

DSS automatically reorders the elements of the matrix alphabetically and makes the first group the intercept. To prevent this, we appended numerals to the beginning of the species name. Macaque is our intercept and the other species are in the order (1) human; (2) chimpanzee; (3) gorilla; (4) orangutan.

Modifications

- Creation of the read counts matrix (named `dss_scores`)
- Creation of the design matrix

**SECTION 4 OF 10: GENERATE THE GLM AND PERFORM THE GATEKEEPING TEST**

This section uses a for loop to independently analyze every site and classify it as “not significantly differential” or “differential”. The classification is based on a chi-squared test of the differences in deviances between the null and fitted models. We save a lot of the parameters and values from DSS and the `glm` function in a results table so that portion of the code will need to be modified extensively.

Design matrix

The `glm` function requires the specification of a design matrix that uses a different format than the one used by DSS. The group names are in the columns and the samples are in the rows. If the sample is in a particular group, then the value is `1`. Otherwise the value is `0`. The group that is used as the intercept is not included in the matrix.

During the loop, the results are stored in four matrices.

- `results_all_sites`
  - This the master output table. It has 54 columns containing parameters from DSS, values from the glm model, read counts, p-values, and other information. The full specification is listed in the comment block at the beginning of the section.
  - The matrix is initially populated with `0` and updated during each iteration of the loop.
- `flag_gateway_test`
  - This is a one column matrix that holds a character flag indicating whether a site is not significantly differential (`GNS`) or differential (`GD`).
  - The matrix is initially populated with `GNS` and updated after the Benjamini-Hochberg correction.
- `flag_change_type`
  - This is a one column matrix that holds a character flag indicating the type of change. The flag is 4 characters long. Each character position represents the comparison of one of the species to macaque. The first position is for human; the second is for chimpanzee; the third is for gorilla; the fourth is for orangutan. There are three types of flags. The first is for sites that are not differential; they have a flag of `ssss`. The second is for sites that

are differential but can't be classified; they have a flag of `OOOO`. The third is for differential sites that can be classified; each class has a different flag. In these flags, each character position has 3 possible values: `s` for same as macaque; `G` for greater than macaque; `L` for less than macaque. For example, a site that is increased only in human will have a flag of `Gsss`, and a site that is decreased in chimpanzee and orangutan will have a flag of `sLsL`.
    - The matrix is initially populated with `ssss` and updated in SECTION 5.
- `constraint_test_pvalues`
    - This is a 15 column matrix that holds the p-values for the 15 constraint tests.
    - The matrix is initially populated with `1` and updated in SECTION 5.

<u>Modifications</u>
- Creation of the design matrix
- Creation of the `results_all_sites` matrix
- Initial values in the `flag_change_type` matrix
- Creation of the `constraint_test_pvalues` matrix
- Column names in the `score_data` matrix used by the `glm` function
- Formula in the `glm` command
- Degrees of freedom in the `pchisq` command
- Values to save in the `results_all_sites` matrix
- Column numbers for the `p.adjust` command (one for raw p-value and one for adjusted p-value)

**SECTION 5 OF 10: PERFORM CONSTRAINT TESTS AND DETERMINE TYPE OF CHANGE**

The contrasts compare the geometric means of the non-macaque species to each other. To determine the coefficients of the constraint matrices, it is helpful to work out the comparisons using equations containing the geometric means. A few things to note:
- Geometric means use multiplication instead of addition.
- The terms on each side of the equation are raised to the power of $1/n$ where $n$ is the number of terms on that side of the equation.
- The coefficients on the left hand side of the equation are entered into the constraint matrix as positive numbers.
- The coefficients on the right hand side of the equation are entered into the constraint matrix as negative numbers.
- Our constraint matrices are in the order: (1) human; (2) chimpanzee; (3) gorilla; (4) orangutan

Example: Human-specific changes. The constraint matrix is [1 -¼ -¼ -¼]

$$\mu_H \neq (\mu_C \mu_G \mu_O \mu_M)^{1/4}$$

$$e^{\beta_M + \beta_H} \neq (e^{\beta_M + \beta_C}\ e^{\beta_M + \beta_G}\ e^{\beta_M + \beta_O}\ e^{\beta_M})^{1/4}$$

$$e^{\beta_M} e^{\beta_H} \neq e^{\beta_M}(e^{\beta_C}\ e^{\beta_G}\ e^{\beta_O})^{1/4}$$

$$e^{\beta_H} \neq (e^{\beta_C}\ e^{\beta_G}\ e^{\beta_O})^{1/4}$$

$$log(e^{\beta_H}) \neq log((e^{\beta_C}\ e^{\beta_G}\ e^{\beta_O})^{1/4})$$

$$\beta_H \neq \frac{\beta_C + \beta_G + \beta_O}{4}$$

Example: Human/Chimpanzee changes. The constraint matrix is [½ ½ -⅓ -⅓]

$$(\mu_H \mu_C)^{1/2} \neq (\mu_G \mu_O \mu_M)^{1/3}$$

$$(e^{\beta_M + \beta_H}\ e^{\beta_M + \beta_C})^{1/2} \neq (e^{\beta_M + \beta_G}\ e^{\beta_M + \beta_O}\ e^{\beta_M})^{1/3}$$

$$e^{\beta_M}(e^{\beta_H}\ e^{\beta_C})^{1/2} \neq e^{\beta_M}(\ e^{\beta_G}\ e^{\beta_O})^{1/3}$$

$$(e^{\beta_H}\ e^{\beta_C})^{1/2} \neq (\ e^{\beta_G}\ e^{\beta_O})^{1/3}$$

$$log((e^{\beta_H}\ e^{\beta_C})^{1/2}) \neq log((\ e^{\beta_G}\ e^{\beta_O})^{1/3})$$

$$\frac{\beta_H + \beta_C}{2} \neq \frac{\beta_G + \beta_O}{3}$$

Modifications

- Creation of the constraint matrices
- Column names in the `score_data` matrix used by the `glm` function
- Formula in the `glm` command
- Calculation of the contrast p-values
- Updating the values in the `constraint_test_pvalues` matrix
- Updating the values in the `flag_change_type` matrix

**SECTION 6 OF 10: UPDATE AND FINALIZE THE "results_all_sites" DATA FRAME**

Update the `results_all_sites` matrix by adding the location information (stored in a data frame called `locations_all_sites`) and three matrices modified during the for loop in section 5:

- `flag_gateway_test`
- `flag_change_type`
- `constraint_test_pvalues`

Modifications

- Creation of the `locations_all_sites` data frame

**SECTION 7 OF 10: CALCULATE TOTALS AND CREATE TABLES CONTAINING SUBSETS OF THE RESULTS**

This section creates subsets for each class of DHS site that are then written to output files in SECTION 8. This section also calculates totals that are used in SECTION 10. This section can be removed completely.

**SECTION 8 OF 10: WRITE RESULTS TO FILES**

There are 67 output files generated in this section. The output file named `glm_analysis.all_sites.all_information.txt` is a master file that has all of the information for all of the DHS sites. There are 2 files for all of the non differential sites and 2 files for all of the differential sites. One of the files has all of the information and has a .txt extension. The other file contains only the location information and has a .bed extension. For each category of differential sites, there are 4 files; 2 files for accessibility decreases (one bed; one txt) and 2 files for accessibility increases (one bed; one txt). These are the same types of files as the ones for the non differential sites. Only the code writing the master file needs to be kept. All of the other code in this section can be removed.

**SECTION 9 OF 10: CALCULATE CHECKSUM AND PRINT STATS**

The checksums are generated to check for errors. All checksum values should be zero. Statistics are generated for summary purposes (e.g. figures and tables). This section can be removed.

**SECTION 10 OF 10: FINISH UP**

Close the connection to the log file. This section does not need to be modified.
