## Supplementary material for "Evaluating chromatin accessibility differences across multiple primate species using a joint modelling approach": supplementary_figures_and_tables.pdf

### FIG. S1

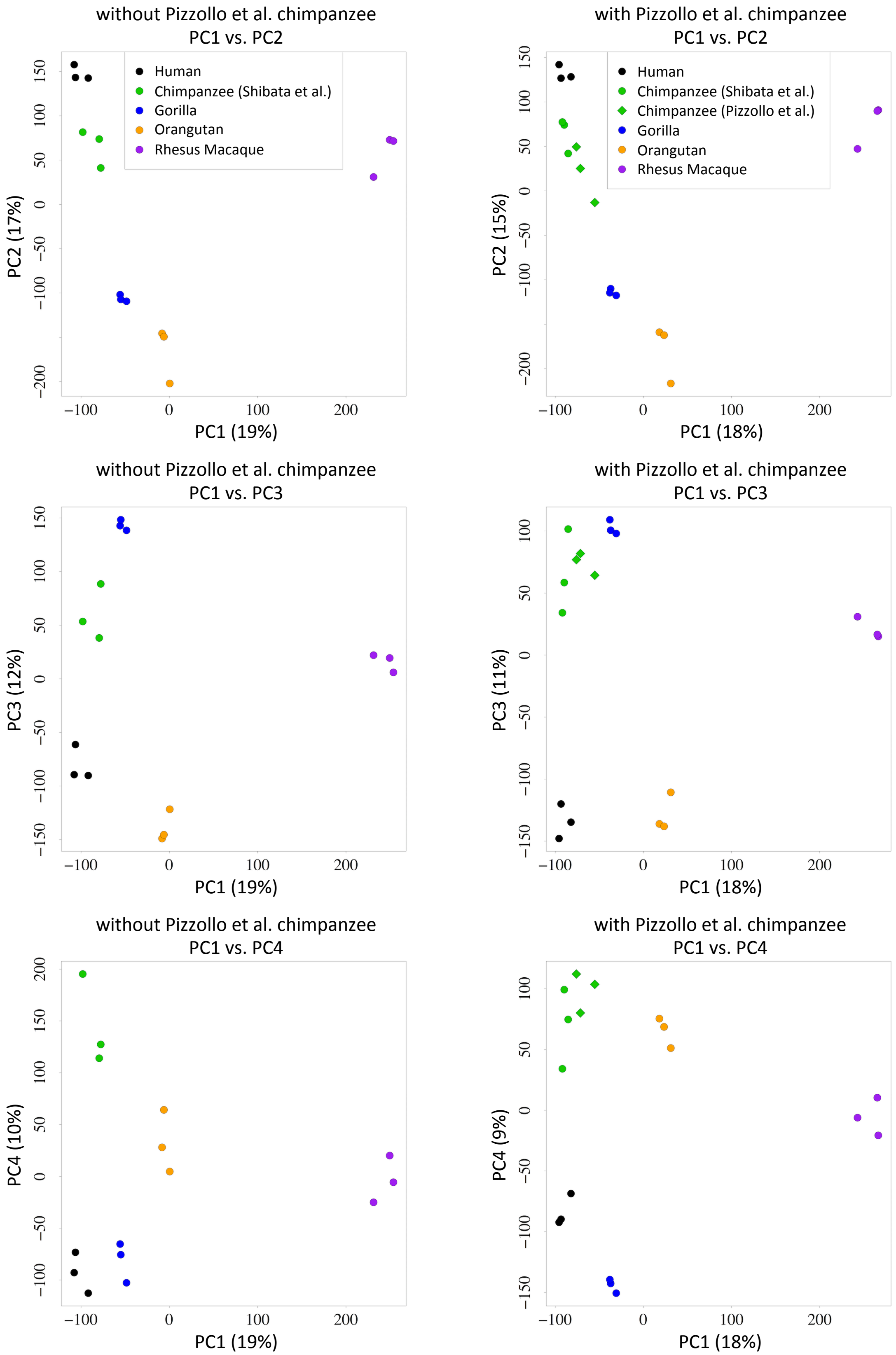

**FIG. S1.** —Principal components analysis of the samples analyzed in this study with (right side) and without (left side) three additional chimpanzee samples from Pizzollo et al. 2018. (Top) Principal component 1 (x-axis) vs. principal component 2 (y-axis). (Middle) Principal component 1 (x-axis) vs. principal component 3 (y-axis). (Bottom) Principal component 1 (x-axis) vs. principal component 4 (y-axis). The original samples are represented by filled circles and the Pizzollo et al. samples are represented by filled diamonds. Human samples are in black, chimpanzee samples are in green, gorilla samples are in blue, orangutan samples are in orange, and rhesus macaque samples are in purple.

### FIG. S2

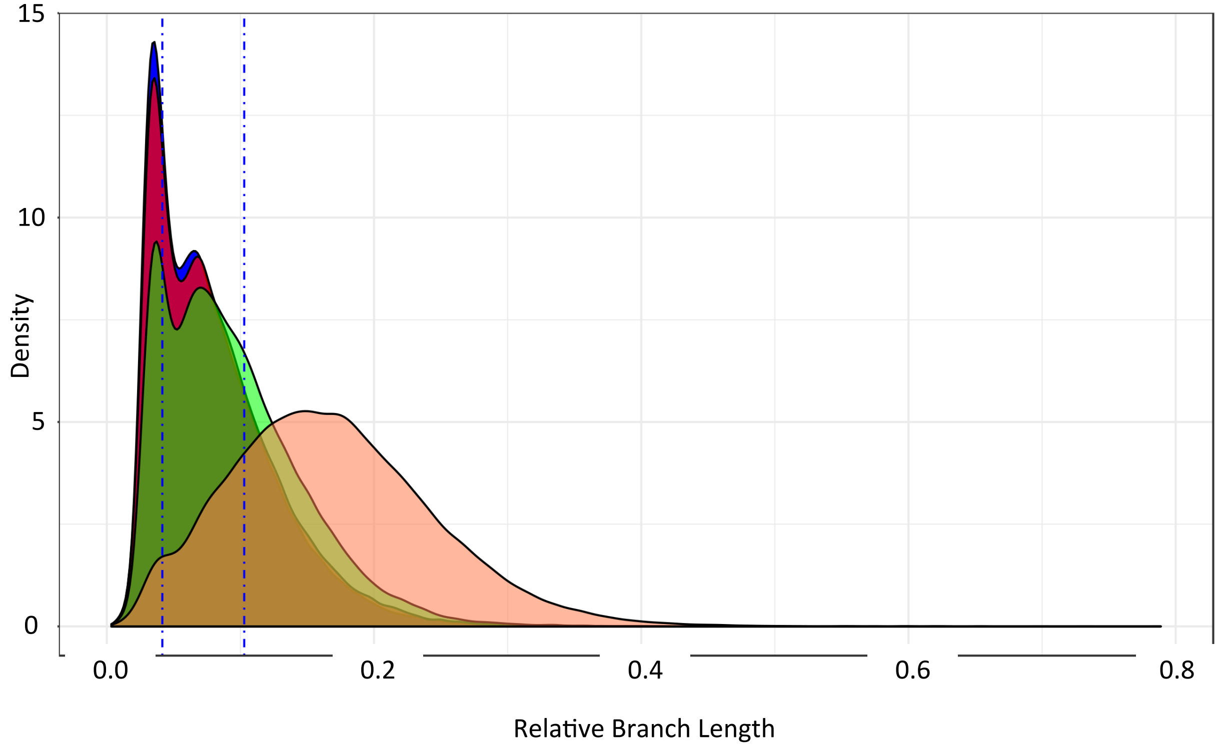

**FIG. S2. —**Distribution of relative branch lengths across a random set of approximately 50,000 genomic regions. The distribution of the relative branch length, which correlates with substitution rate, for human (blue), chimpanzee (red), gorilla (green), and orangutan (coral). The vertical blue lines represent the 1st and 3rd quartile for the human branches. Adapted from Berrio et al 2019.

### FIG. S3

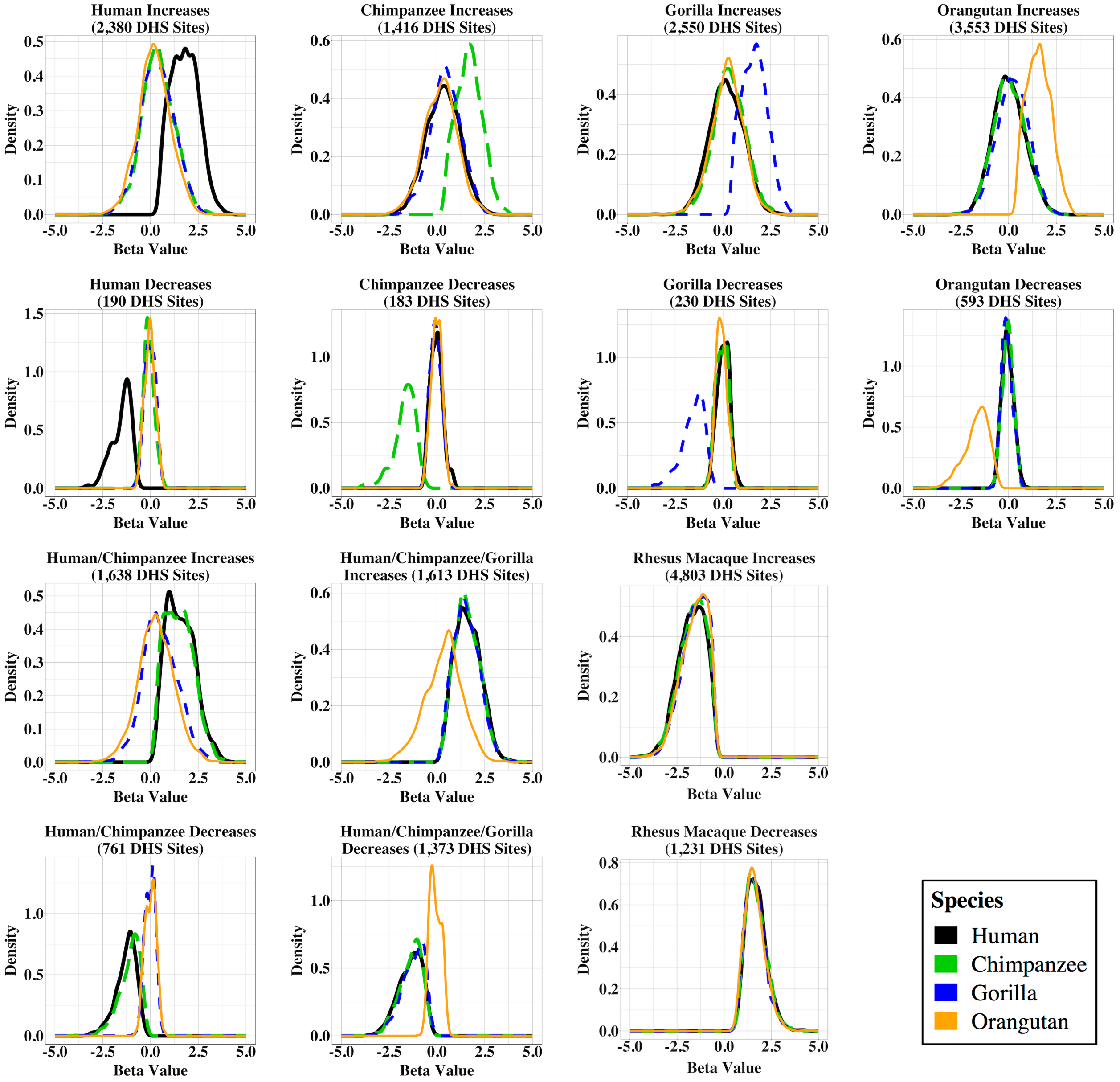

**FIG. S3.** —Distribution of beta values for differential chromatin accessibility due to a single change. Density plots showing the beta values for human (black), chimpanzee (green), gorilla (blue), and orangutan (orange). Each plot is for a different category of differential site.

### FIG. S4

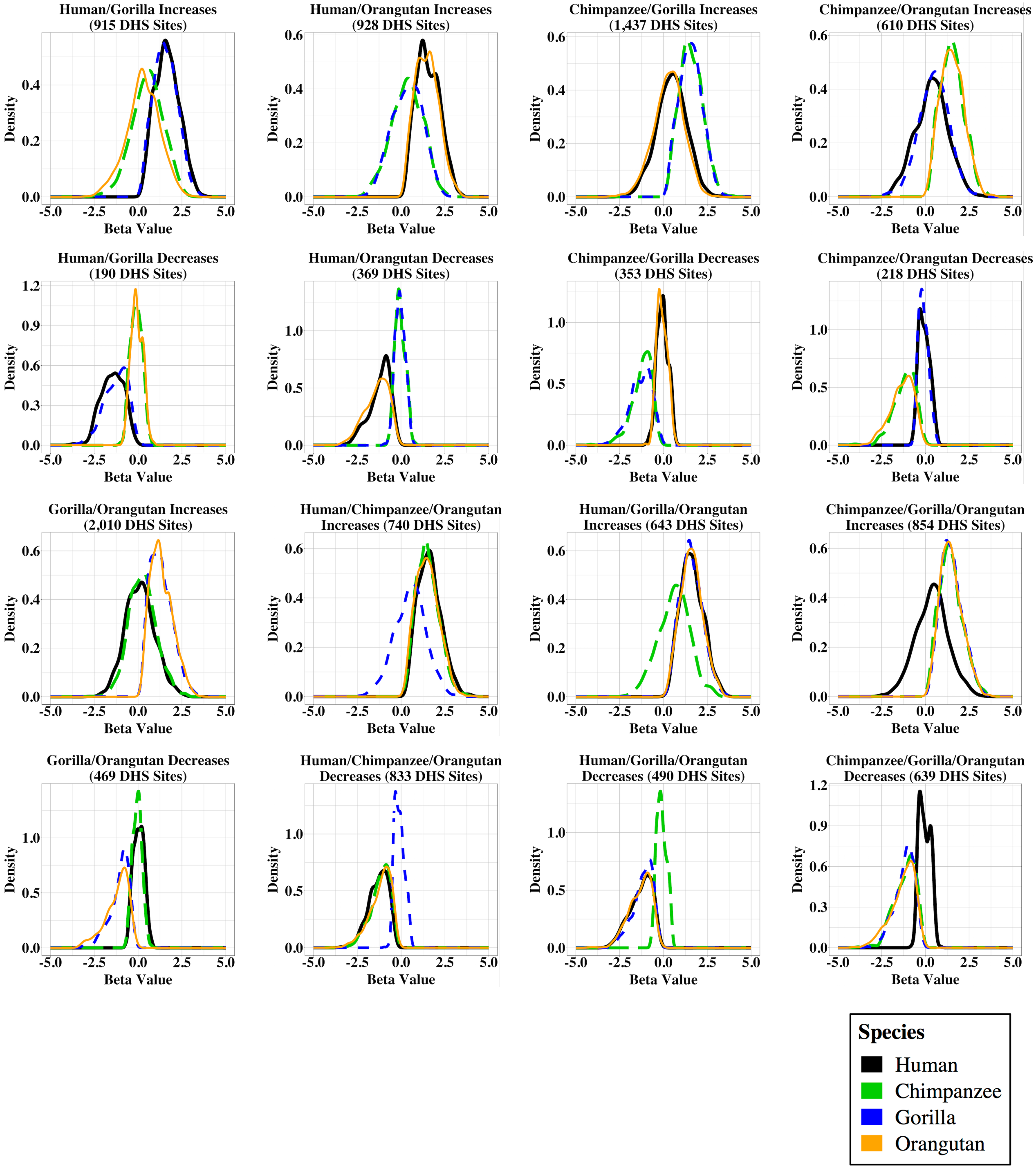

**FIG. S4. —**Distribution of beta values for differential chromatin accessibility due to multiple changes. Density plots showing the beta values for human (black), chimpanzee (green), gorilla (blue), and orangutan (orange). Each plot is for a different category of differential site.

### FIG. S5

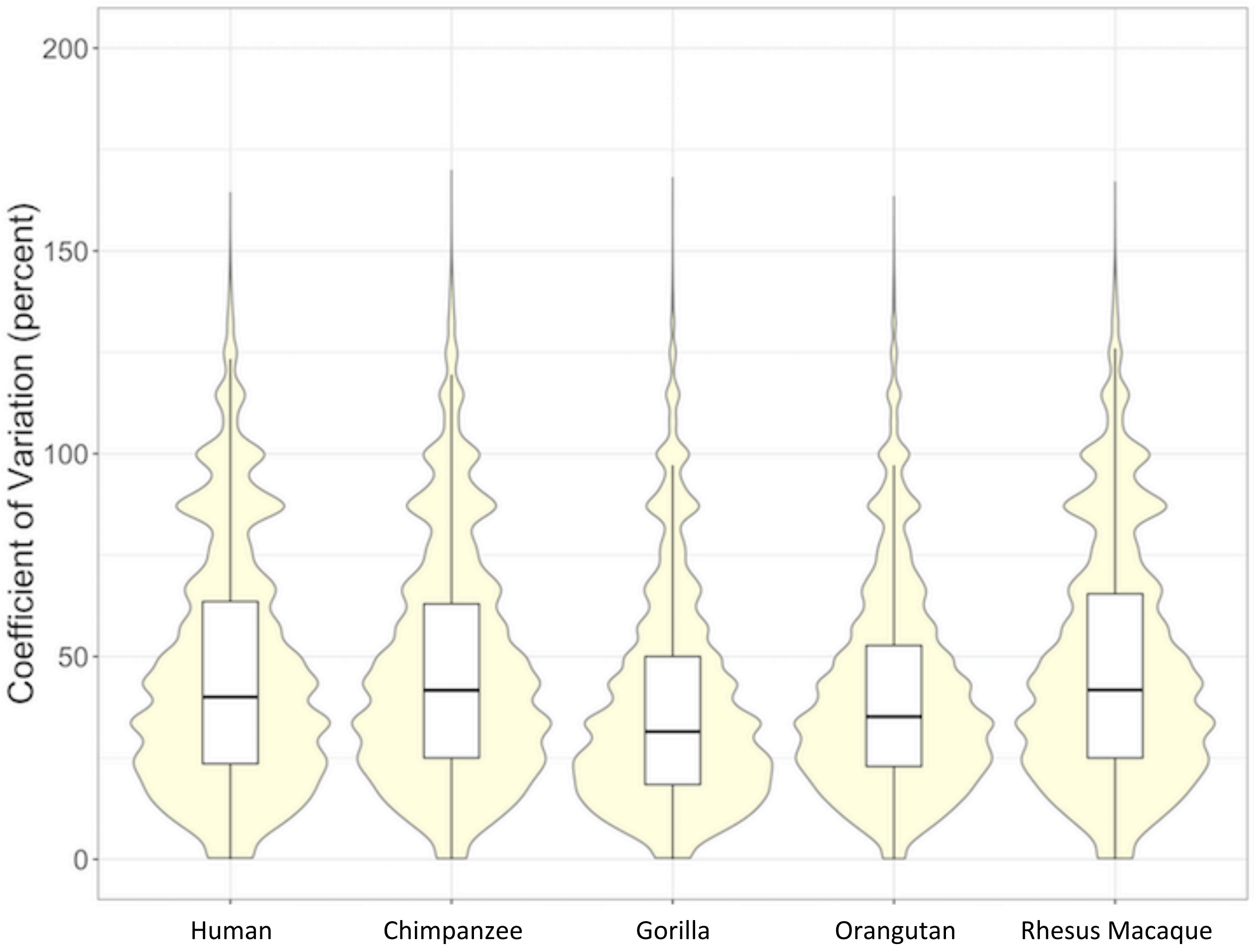

**FIG. S5.** —Violin plots showing distribution of intra-species variation in normalized read counts calculated for each DHS site.

### FIG. S6

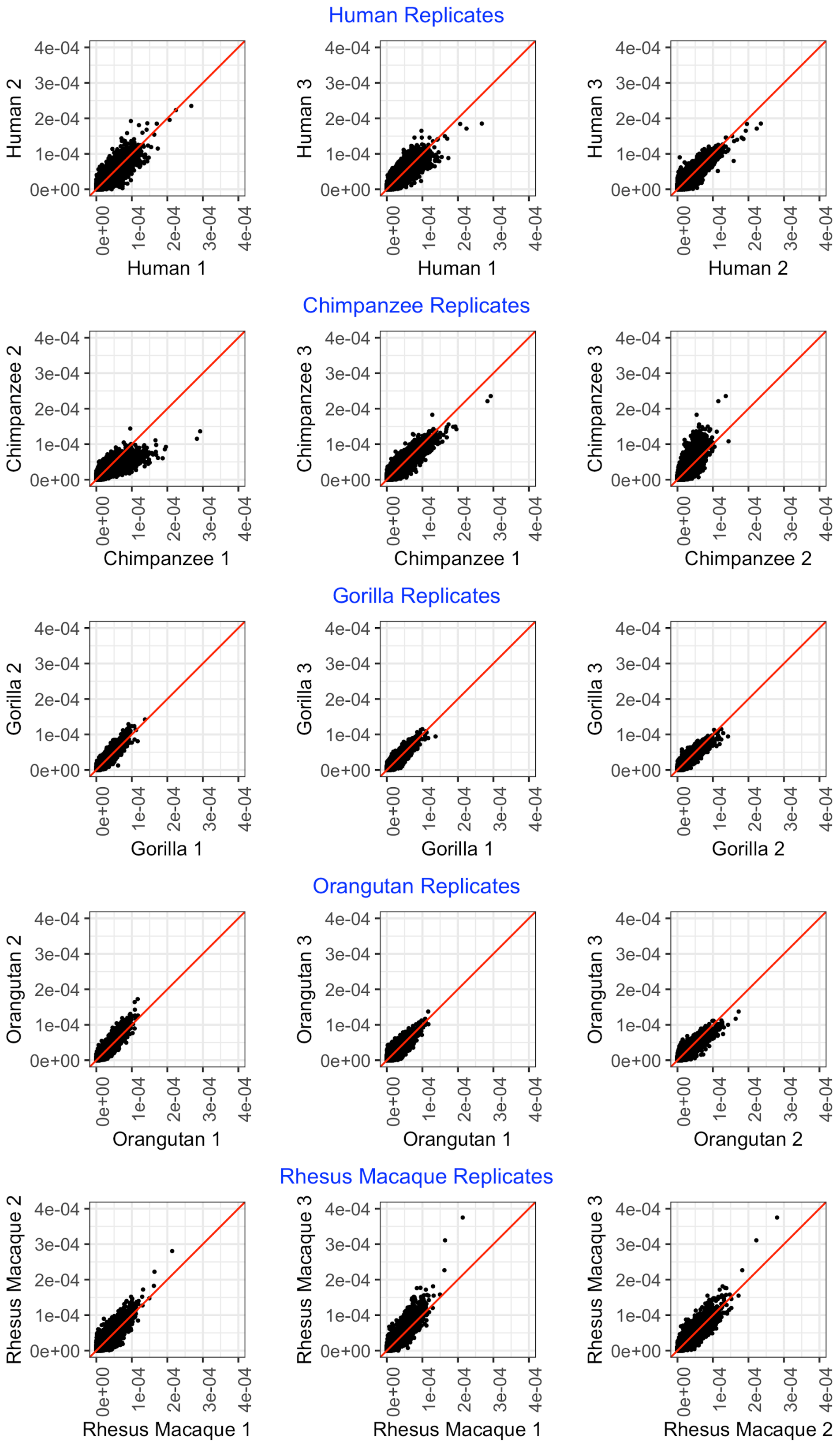

**FIG. S6. —**Scatter plots showing differences in normalized read counts between biological replicates calculated for each DHS site. (Left) Replicate 1 (x-axis) vs. replicate 2 (y-axis). (Middle) Replicate 1 (x-axis) vs. replicate 3 (y-axis). (Right) Replicate 2 (x-axis) vs. replicate 3 (y-axis).

### FIG. S7

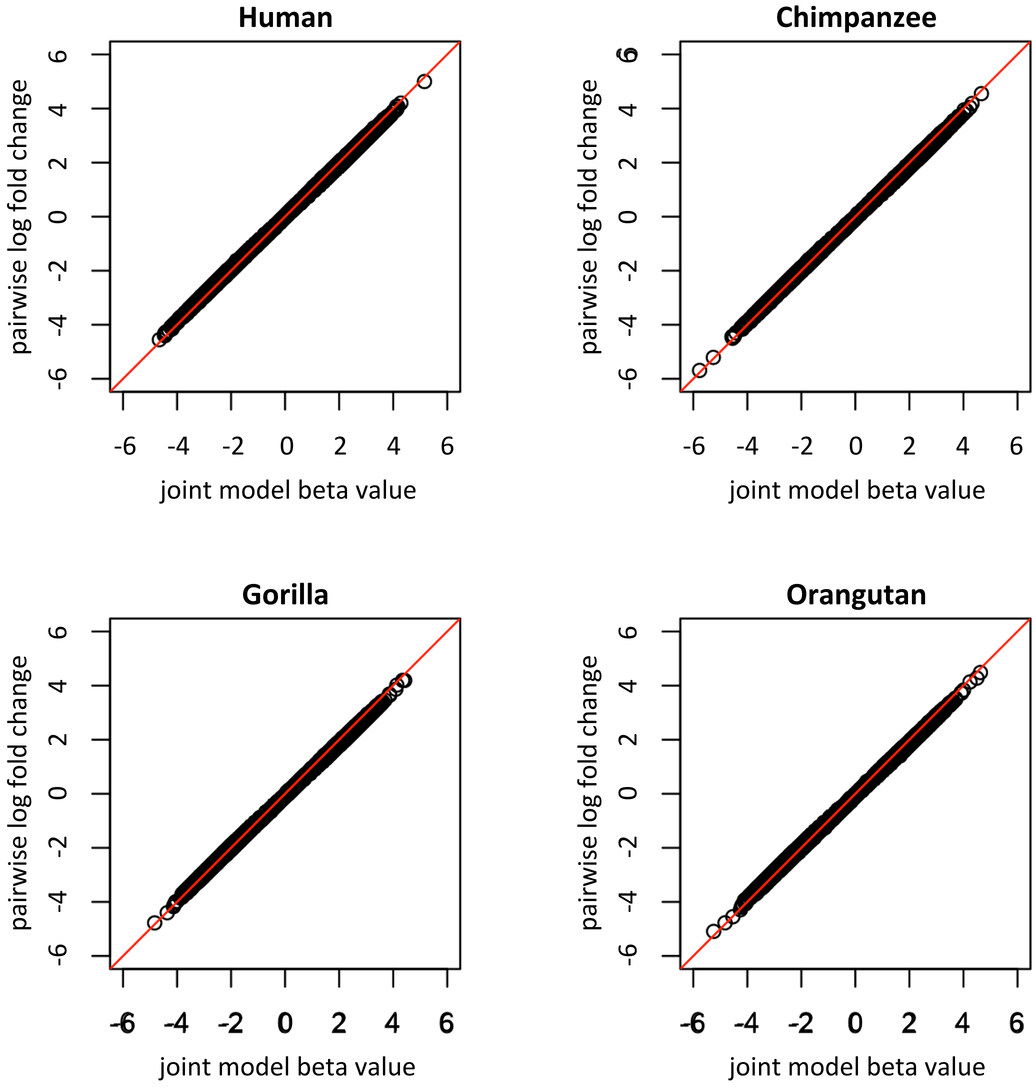

**FIG. S7. —**Comparison of pairwise log fold change values and joint model beta values. Joint model beta values (x-axis) and pairwise log fold change values (y-axis) are shown on the natural log scale. Values are from all DHS sites.

### TABLE S1

| Sample | Coriell ID | Technical Replicates | Library Date | Number of Sequencing Lanes | Sequencer Type (Machine Name) | Biopsy Source | Sex | Age at Sampling | Notes |
|---|---|---|---|---|---|---|---|---|---|
| Human 1 | AG16409 | 1 | 8/6/08 | 4 (2 lanes per sequencer) | Genome Analyzer II (HWI-EAS215) (HWUSI-EAS465) | Unknown | Male | 12 YR | Caucasian |
| Human 2 | GM02185 | 1 | 12/16/08 | 4 | Genome Analyzer II (HWUSI-EAS465) | Unknown | Male | 36 YR | Caucasian |
| Human 3 | GM05879 | 1 | 12/16/08 | 4 | Genome Analyzer II (HWUSI-EAS465) | Arm | Female | 48 YR | Caucasian |
| Chimpanzee 1 | S003649 | 1 | 8/6/08 | 4 (2 lanes per sequencer) | Genome Analyzer II (HWI-EAS215) (HWUSI-EAS465) | Unknown | Male | 9 YR | from the Yerkes Primates name is Josh |
| Chimpanzee 2 | S003624 | 1 | 11/21/08 | 3 (2 lanes in one run; 1 lane in another) | Genome Analyzer IIx (HWUSI-EAS1676) | Unknown | Male | 14 YR | from the Yerkes Primates name is Tank |
| Chimpanzee 3 | S007603 | 1 | 11/21/08 | 4 | Genome Analyzer II (HWUSI-EAS465) | Unknown | Male | 16 YR | from the Yerkes Primates name is Donald |
| Chimpanzee 4 | PR00238 | 1 | 9/13/12 | 1 | HiSeq 2000 (D3NH4HQ1) | Unknown | Male | 24 YR | name is Clint |
| Chimpanzee 5 | S006007 | 1 | 9/13/12 | 1 | HiSeq 2000 (D3NH4HQ1) | Unknown | Male | 22 YR | |
| Chimpanzee 6 | S008975 | 1 | 9/13/12 | 1 | HiSeq 2000 (D3NH4HQ1) | Unknown | Male | 19 YR | |
| Gorilla 1 | PR00107 | 1 | 10/17/12 | 1 | HiSeq 2000 (HWI-ST1293) | Unknown | Male | 19 YR | |
| Gorilla 2 | PR00669 | 1 | 10/17/12 | 1 | HiSeq 2000 (HWI-ST1293) | Unknown | Male | 15 YR | |
| Gorilla 3 | PR01103 | 1 | 10/17/12 | 1 | HiSeq 2000 (HWI-ST1293) | Unknown | Male | 16 YR | |
| Orangutan 1 | PR00054 | 1 | 8/3/12 | 1 | HiSeq 2000 (D3NH4HQ1) | Unknown | Male | 4 YR | Sumatran orangutan (Pongo pygmaeus) |
| Orangutan 2 | PR01107 | 1 | 8/3/12 | 1 | HiSeq 2000 (D3NH4HQ1) | Unknown | Male | 11 YR | Sumatran orangutan (Pongo pygmaeus) |
| Orangutan 3 | GM04272 | 1 | 8/24/12 | 1 | HiSeq 2000 (D3NH4HQ1) | Unknown | Male | 10 YR | Sumatran orangutan (Pongo pygmaeus) |
| Rhesus Macaque 1 | AG06252 | 2 | 8/6/08 | 6 (3 lanes per technical replicate) | Genome Analyzer IIx (HWUSI-EAS407) | Arm | Female | 5 YR | Indian strain |
| Rhesus Macaque 2 | AG08305 | 2 | 12/16/08 | 7 total<br>*Technical replicate #1* 4 lanes (3 lanes in one run, 1 lane in another)<br>*Technical replicate #2* 3 lanes (2 lanes in one run, 1 lane in another) | *Technical replicate #1* Genome Analyzer II (HWUSI-EAS465)<br>*Technical replicate #2* Genome Analyzer IIx (HWUSI-EAS1676) for 1 lane Genome Analyzer II (HWI-EAS215) for 2 lanes | Arm | Male | 1 YR | |
| Rhesus Macaque 3 | AG08308 | 1 | 11/21/08 | 4 (3 lanes in one run; 1 lane in another) | Genome Analyzer II (unknown) | Arm | Male | 1 YR | |

Sample information

### TABLE S2

| Replicate | Raw reads | Tier 1 (No mismatches) Unique hits | | Multiple hits | | Unaligned | | Tier 2 (1 mismatch) Unique hits | | Multiple hits | | Unaligned | |
|---|---|---|---|---|---|---|---|---|---|---|---|---|---|
| Human 1 | 50,938,241 | 32,879,743 | 65% | 12,283,795 | 24% | 5,774,703 | 11% | 2,136,098 | 4% | 2,347,034 | 5% | 1,291,571 | 3% |
| Human 2 | 33,630,278 | 22,180,616 | 66% | 8,140,672 | 24% | 3,308,990 | 10% | 1,141,379 | 3% | 1,654,187 | 5% | 513,424 | 2% |
| Human 3 | 23,844,246 | 15,548,666 | 65% | 5,556,547 | 23% | 2,739,033 | 11% | 1,017,236 | 4% | 1,263,353 | 5% | 458,444 | 2% |
| Chimpanzee 1 | 50,096,931 | 31,674,269 | 63% | 13,476,106 | 27% | 4,946,556 | 10% | 1,612,581 | 3% | 1,735,149 | 3% | 1,598,826 | 3% |
| Chimpanzee 2 | 111,972,567 | 72,746,554 | 65% | 29,841,886 | 27% | 9,384,127 | 8% | 2,750,866 | 2% | 3,307,427 | 3% | 3,325,834 | 3% |
| Chimpanzee 3 | 34,388,687 | 21,848,911 | 64% | 9,120,646 | 27% | 3,419,130 | 10% | 1,169,139 | 3% | 1,379,792 | 4% | 870,199 | 3% |
| Gorilla 1 | 200,394,049 | 113,954,996 | 57% | 46,452,406 | 23% | 39,986,647 | 20% | 17,164,167 | 9% | 15,177,562 | 8% | 7,644,918 | 4% |
| Gorilla 2 | 207,690,245 | 116,195,086 | 56% | 46,244,395 | 22% | 45,250,764 | 22% | 18,247,477 | 9% | 15,755,405 | 8% | 11,247,882 | 5% |
| Gorilla 3 | 218,321,295 | 139,735,317 | 64% | 56,153,448 | 26% | 22,432,530 | 10% | 7,057,645 | 3% | 8,422,152 | 4% | 6,952,733 | 3% |
| Orangutan 1 | 204,329,161 | 126,205,138 | 62% | 44,601,400 | 22% | 33,522,623 | 16% | 9,849,034 | 5% | 10,301,994 | 5% | 13,371,595 | 7% |
| Orangutan 2 | 223,997,352 | 136,190,707 | 61% | 51,142,667 | 23% | 36,663,978 | 16% | 10,465,410 | 5% | 12,391,751 | 6% | 13,806,817 | 6% |
| Orangutan 3 | 228,118,747 | 124,383,888 | 55% | 46,679,518 | 20% | 57,055,341 | 25% | 23,616,339 | 10% | 20,069,852 | 9% | 13,369,150 | 6% |
| Rhesus Macaque 1 | 70,897,729 | 34,129,807 | 48% | 12,911,828 | 18% | 23,856,094 | 34% | 9,092,082 | 13% | 11,503,154 | 16% | 3,260,858 | 5% |
| Rhesus Macaque 2 | 69,917,208 | 42,438,629 | 61% | 14,314,693 | 20% | 13,163,886 | 19% | 4,609,783 | 7% | 4,988,172 | 7% | 3,565,931 | 5% |
| Rhesus Macaque 3 | 29,133,743 | 18,596,481 | 64% | 6,046,944 | 21% | 4,490,318 | 15% | 1,456,525 | 5% | 1,666,868 | 6% | 1,366,925 | 5% |

| Replicate | Final mapping results Unique hits | | Multiple hits | | Unaligned | | Final results Unique monoclonal | |
|---|---|---|---|---|---|---|---|---|
| Human 1 | 35,015,841 | 69% | 14,630,829 | 29% | 1,291,571 | 3% | 14,476,479 | 28% |
| Human 2 | 23,321,995 | 69% | 9,794,859 | 29% | 513,424 | 2% | 15,287,873 | 45% |
| Human 3 | 16,565,902 | 69% | 6,819,900 | 29% | 458,444 | 2% | 13,516,977 | 57% |
| Chimpanzee 1 | 33,286,850 | 66% | 15,211,255 | 30% | 1,598,826 | 3% | 11,245,529 | 22% |
| Chimpanzee 2 | 75,497,420 | 67% | 33,149,313 | 30% | 3,325,834 | 3% | 51,812,496 | 46% |
| Chimpanzee 3 | 23,018,050 | 67% | 10,500,438 | 31% | 870,199 | 3% | 15,660,736 | 46% |
| Gorilla 1 | 131,119,163 | 65% | 61,629,968 | 31% | 7,644,918 | 4% | 108,696,016 | 54% |
| Gorilla 2 | 134,442,563 | 65% | 61,999,800 | 30% | 11,247,882 | 5% | 95,844,959 | 46% |
| Gorilla 3 | 146,792,962 | 67% | 64,575,600 | 30% | 6,952,733 | 3% | 119,415,564 | 55% |
| Orangutan 1 | 136,054,172 | 67% | 54,903,394 | 27% | 13,371,595 | 7% | 99,830,828 | 49% |
| Orangutan 2 | 146,656,117 | 65% | 63,534,418 | 28% | 13,806,817 | 6% | 47,027,229 | 21% |
| Orangutan 3 | 148,000,227 | 65% | 66,749,370 | 29% | 13,369,150 | 6% | 108,387,926 | 48% |
| Rhesus Macaque 1 | 43,221,889 | 61% | 24,414,982 | 34% | 3,260,858 | 5% | 40,813,669 | 58% |
| Rhesus Macaque 2 | 47,048,412 | 67% | 19,302,865 | 28% | 3,565,931 | 5% | 23,197,615 | 33% |
| Rhesus Macaque 3 | 20,053,006 | 69% | 7,713,812 | 26% | 1,366,925 | 5% | 13,836,905 | 47% |

| Replicate | Mapped | After lift over (a) | | Final counts (b) |
|---|---|---|---|---|
| Human 1 | 14,388,372 | 14,388,372 | 100% | 14,014,031 |
| Human 2 | 15,204,571 | 15,204,571 | 100% | 14,810,459 |
| Human 3 | 13,437,729 | 13,437,729 | 100% | 13,105,259 |
| Chimpanzee 1 | 11,245,529 | 10,861,657 | 97% | 10,624,634 |
| Chimpanzee 2 | 47,421,354 | 45,614,631 | 96% | 44,529,183 |
| Chimpanzee 3 | 15,660,736 | 15,129,473 | 97% | 14,783,843 |
| Gorilla 1 | 104,413,179 | 101,111,374 | 97% | 98,813,074 |
| Gorilla 2 | 91,500,123 | 88,494,541 | 97% | 86,567,493 |
| Gorilla 3 | 114,551,049 | 110,815,063 | 97% | 108,405,287 |
| Orangutan 1 | 99,830,828 | 89,033,545 | 89% | 86,998,499 |
| Orangutan 2 | 47,027,229 | 43,114,002 | 92% | 42,076,972 |
| Orangutan 3 | 108,387,926 | 98,211,154 | 91% | 95,803,771 |
| Rhesus Macaque 1 | 40,813,669 | 35,868,548 | 88% | 34,430,825 |
| Rhesus Macaque 2 | 23,197,615 | 20,611,242 | 89% | 20,140,263 |
| Rhesus Macaque 3 | 13,836,905 | 12,332,942 | 89% | 12,040,473 |

Mapping and LiftOver statistics
NOTE. —(a) Human replicates were not lifted over; the numbers are repeated for consistency. Percentages of “After lift over” reads are of mapped reads. (b) Final counts are after reads mapped to sex chromosomes are removed.

### TABLE S3

| Sample | DHS sites |
|---|---|
| Human 1 | 99,030 |
| Human 2 | 84,421 |
| Human 3 | 90,871 |
| Chimpanzee 1 | 163,806 |
| Chimpanzee 2 | 113,653 |
| Chimpanzee 3 | 104,601 |
| Gorilla 1 | 107,869 |
| Gorilla 2 | 122,317 |
| Gorilla 3 | 86,317 |
| Orangutan 1 | 161,994 |
| Orangutan 2 | 180,239 |
| Orangutan 3 | 113,979 |
| Rhesus Macaque 1 | 119,191 |
| Rhesus Macaque 2 | 123,237 |
| Rhesus Macaque 3 | 85,305 |

| Species | DHS sites |
|---|---|
| Human | 62,017 |
| Chimpanzee | 75,700 |
| Gorilla | 63,512 |
| Orangutan | 94,883 |
| Rhesus Macaque | 68,720 |

| Step | DHS Sites |
|---|---|
| Before filtering | 166,331 |
| After filtering for genomic coverage | 124,900 |
| After filtering for zero-count | 89,744 |

DHS site counts at different steps in the analysis pipeline

### TABLE S4

| | Human Chimpanzee Gorilla Orangutan |
|---|---|
| Human | [1 -¼ -¼ -¼] |
| Chimpanzee | [-¼ 1 -¼ -¼] |
| Gorilla | [-¼ -¼ 1 -¼] |
| Orangutan | [-¼ -¼ -¼ 1] |
| Rhesus Macaque | [-¼ -¼ -¼ -¼] |
| Human/Chimpanzee | [½ ½ -⅓ -⅓] |
| Human/Gorilla | [½ -⅓ ½ -⅓] |
| Human/Orangutan | [½ -⅓ -⅓ ½] |
| Chimpanzee/Gorilla | [-⅓ ½ ½ -⅓] |
| Chimpanzee/Orangutan | [-⅓ ½ -⅓ ½] |
| Gorilla/Orangutan | [-⅓ -⅓ ½ ½] |
| Human/Chimpanzee/Gorilla | [⅓ ⅓ ⅓ -½] |
| Human/Chimpanzee/Orangutan | [⅓ ⅓ -½ ⅓] |
| Human/Gorilla/Orangutan | [⅓ -½ ⅓ ⅓] |
| Chimpanzee/Gorilla/Orangutan | [-½ ⅓ ⅓ ⅓] |

Constraint matrices

### TABLE S5

| Class of DHS sites | Foreground 1 | Foreground 2 |
|---|---|---|
| Human increased accessibility | Human | Chimpanzee |
| Human increased accessibility | Human | Gorilla |
| Human decreased accessibility | Human | Chimpanzee |
| Human decreased accessibility | Human | Gorilla |
| Chimpanzee increased accessibility | Chimpanzee | Human |
| Chimpanzee increased accessibility | Chimpanzee | Gorilla |
| Chimpanzee decreased accessibility | Chimpanzee | Human |
| Chimpanzee decreased accessibility | Chimpanzee | Gorilla |
| Gorilla increased accessibility | Gorilla | Human |
| Gorilla increased accessibility | Gorilla | Chimpanzee |
| Gorilla decreased accessibility | Gorilla | Human |
| Gorilla decreased accessibility | Gorilla | Chimpanzee |
| Non differential | Human | Chimpanzee |
| Non differential | Human | Gorilla |
| Non differential | Chimpanzee | Gorilla |

Classes of DHS sites and foregrounds used for comparisons of enrichment of positive selection

### TABLE S6

| |
|---|
| Human increased accessibility |
| Chimpanzee increased accessibility |
| Gorilla increased accessibility |
| Orangutan increased accessibility |
| Human-Chimpanzee increased accessibility |
| Human-Chimpanzee-Gorilla increased accessibility |
| Human decreased accessibility |
| Chimpanzee decreased accessibility |
| Gorilla decreased accessibility |
| Orangutan decreased accessibility |
| Human-Chimpanzee decreased accessibility |
| Human-Chimpanzee-Gorilla decreased accessibility |

Classes of DHS sites used for comparison of the distributions of human zeta values

### TABLE S7

| |
|---|
| Human increased accessibility |
| Chimpanzee increased accessibility |
| Gorilla increased accessibility |
| Orangutan increased accessibility |
| Human-Chimpanzee increased accessibility |
| Human-Chimpanzee-Gorilla increased accessibility |
| Human decreased accessibility |
| Chimpanzee decreased accessibility |
| Gorilla decreased accessibility |
| Orangutan decreased accessibility |
| Human-Chimpanzee decreased accessibility |
| Human-Chimpanzee-Gorilla decreased accessibility |
| Non differential |

Classes of DHS sites used for comparisons of the distributions of cell type specificity values

### TABLE S8

| HUMAN AND CHIMPANZEE CHANGES | | | | | | | | | |
|---|---|---|---|---|---|---|---|---|---|
| | **TOTAL CONSISTENT** | | **Consistent (single species)** | | **Consistent (multi-species)** | | **Not consistent** | | **Total** |
| Human increases | **336** | **98%** | 245 | 72% | 91 | 27% | 6 | 2% | 342 |
| Chimpanzee increases | **218** | **93%** | 104 | 44% | 114 | 49% | 16 | 7% | 234 |
| Human decreases | **128** | **86%** | 21 | 14% | 107 | 72% | 18 | 12% | 148 |
| Chimpanzee decreases | **85** | **87%** | 17 | 17% | 68 | 69% | 13 | 13% | 98 |

| COMMON SITES | | | | | | | | |
|---|---|---|---|---|---|---|---|---|
| **TOTAL CONSISTENT** | | **Non differential** | | **Gorilla and/or orangutan** | | **Not consistent** | | **Total** |
| **1,099** | **95%** | 1,043 | 90% | 56 | 5% | 55 | 5% | 1,154 |

Results from comparison to earlier study (Shibata et al., 2012, PLoS Genetics)
NOTE. —Increase refers to an increase in chromatin accessibility compared to rhesus macaque. Decrease refers to a decrease in chromatin accessibility compared to rhesus macaque. Total refers to the number of overlapped sites. Numbers in consistent and not consistent box refer to calls made by the generalized linear model.

### TABLE S9

| Step | Type of Change | Shibata et al. Pipeline | This Pipeline |
|---|---|---|---|
| Mapping | Mapping software | BWA | Bowtie |
| Mapping | Mapping parameters | Up to 3 mismatches | Up to 1 mismatch |
| Mapping | Mapping parameters | Multiple locations allowed | Unique locations only |
| Mapping | Chimpanzee genome | panTro2 | panTro4 |
| Mapping | Rhesus Macaque genome | rheMac2 | rheMac3 |
| DHS site identification | Peak calling program | F-Seq | MACS2 |
| DHS site identification | Per-species DHS sites | Union set of top 100,000 peaks present in any replicate | Union set of all peaks present in at least two replicates |
| DHS site identification | DHS site width | DHS sites less than 300 bases were extended to 300 bases | DHS site widths unchanged |
| DHS site identification | Windowing | DHS sites were divided into overlapping windows of 300 bases | None |
| DHS site filtering | Genomic coverage | None | Genomic coverage of at least 95% of the bases in every species |
| DHS site filtering | Zero-count filtering | At least one replicate had to have non-zero counts | At least two replicates had to have non-zero counts |

| | Original sites | With overlapping reads | | With 1+ replicate peaks | | With 2+ replicate peaks | | Passed genomic coverage filter | | Passed zero-count filter and analyzed | | Not analyzed | |
|---|---|---|---|---|---|---|---|---|---|---|---|---|---|
| Human Increases | 836 | 831 | 99% | 761 | 91% | 606 | 72% | 531 | 64% | **310** | **37%** | 526 | 63% |
| Chimpanzee Increases | 676 | 650 | 96% | 562 | 83% | 442 | 65% | 399 | 59% | **215** | **32%** | 461 | 68% |
| Human Decreases | 286 | 281 | 98% | N/A | N/A | N/A | N/A | 198 | 69% | **133** | **47%** | 153 | 53% |
| Chimpanzee Decreases | 211 | 194 | 92% | N/A | N/A | N/A | N/A | 135 | 64% | **89** | **42%** | 122 | 58% |

Differences in analysis pipeline compared to earlier study (Shibata et al., 2012, PLoS Genetics)

### TABLE S10

**INCREASED ACCESSIBILITY COMPARED TO RHESUS MACAQUE**

| | Human | | Chimpanzee | | Gorilla | | Orangutan | | H-C | | H-C-G | |
|---|---|---|---|---|---|---|---|---|---|---|---|---|
| **Proximal** | 60 | 3% | 53 | 4% | 221 | 9% | 344 | 10% | 43 | 3% | 82 | 5% |
| **Distal** | 1,975 | 83% | 911 | 64% | 1,515 | 59% | 1,811 | 51% | 1,354 | 83% | 1,256 | 78% |
| **Unannotated** | 345 | 14% | 452 | 32% | 814 | 32% | 1,398 | 39% | 241 | 15% | 275 | 17% |
| **Total** | **2,380** | | **1,416** | | **2,550** | | **3,553** | | **1,638** | | **1,613** | |

**DECREASED ACCESSIBILITY COMPARED TO RHESUS MACAQUE**

| | Human | | Chimpanzee | | Gorilla | | Orangutan | | H-C | | H-C-G | |
|---|---|---|---|---|---|---|---|---|---|---|---|---|
| **Proximal** | 37 | 19% | 41 | 22% | 27 | 12% | 124 | 21% | 264 | 35% | 145 | 11% |
| **Distal** | 90 | 47% | 115 | 63% | 165 | 72% | 356 | 60% | 273 | 36% | 751 | 55% |
| **Unannotated** | 63 | 33% | 27 | 15% | 38 | 17% | 113 | 19% | 224 | 29% | 477 | 35% |
| **Total** | **190** | | **183** | | **230** | | **593** | | **761** | | **1,373** | |

| | Non Differential | |
|---|---|---|
| **Proximal** | 11,850 | 22% |
| **Distal** | 30,371 | 57% |
| **Unannotated** | 10,857 | 20% |
| **Total** | **53,078** | |

Location of DHS sites

NOTE. —Percentages are of the total number of DHS sites. H-C: human-chimpanzee internal branch. H-C-G: human-chimpanzee-gorilla internal branch.
